# Targeting Mitochondrial Dysfunction with Mdivi-1 Confers Therapeutic Protection in Cell Culture and Mouse Models of Mustard Keratopathy

**DOI:** 10.64898/2026.08.27.742467

**Authors:** Arpan G. Mazumder, Evan Magarychoff, Hamid Alemi, Rahul Raghav, Michael T. Chin, Zhen Yan, Christopher D. Wiley, M. Elizabeth Fini

## Abstract

Mustard keratopathy, caused by exposure of the cornea to chemical warfare vesicants sulfur or nitrogen mustard, can lead to severe and often irreversible vision loss. Despite considerable efforts to develop medical countermeasures, including anti-inflammatory, antioxidant, anti-fibrotic, and anti-angiogenic therapies, no treatment effectively targets the underlying mechanisms responsible for mustard-induced tissue injury or prevents long-term disease recurrence. In the present study, we comprehensively define mitochondrial mechanisms underlying nitrogen mustard-induced corneal injury in both a human cell culture model in vitro and a mouse model in vivo. We demonstrate that nitrogen mustard rapidly induces excessive mitochondrial fragmentation, bioenergetic collapse, membrane depolarization, oxidative stress, intracellular acidification, mitophagy, and apoptotic cell death. Pharmacological intervention with the small molecule Mdivi-1 preserves mitochondrial structure and function, restores cellular metabolism, reduces oxidative damage, and markedly improves corneal epithelial integrity and tissue repair following nitrogen mustard exposure. Mdivi-1 targets DNM1L (aka Drp1), a mitochondria-localized dynamin-related GTPase that has been shown to execute mitochondrial fission and facilitate the autophagic elimination of damaged mitochondrial components, and our findings are consistent with this idea. Collectively, these findings establish mitochondrial dysfunction as a central pathological mechanism in mustard keratopathy and identify DNM1L-mediated mitochondrial remodeling as a therapeutically actionable target. The findings provide early preclinical evidence supporting mitochondrial-directed therapy as a promising medical countermeasure for treatment of mustard keratopathy.

**One Sentence Summary:** The study findings establish mitochondrial dysfunction as a central pathological mechanism in mustard keratopathy and identify DNM1L-mediated mitochondrial remodeling as a therapeutically actionable target.

## 1. Introduction

A high-consequence chemical emergency is a major public health concern demanding development of critical medical countermeasures against chemical threats for health security and public preparedness. Mustard keratopathy, caused by exposure of the cornea to sulfur or nitrogen mustard vesicants, chemical warfare agents, can lead to severe and often irreversible vision loss [1]. Ocular injury follows a characteristic biphasic course, with an acute phase of epithelial damage and inflammation. The necroptosis programmed cell death pathway is reported to be dominant over apoptosis as a driver of acute corneal epithelial damage [2]. Unlike apoptosis, necroptosis is a highly lytic and pro-inflammatory form of cell death. Nitrogen mustard also triggers ferroptosis in corneal stromal cells [3]. This leads to an overwhelming burst in lipid peroxidation and reactive oxygen species generation, which further fuels the local inflammatory environment. The acute phase is followed by a prolonged latent period that can culminate in recurrent or delayed-onset pathology years after the initial exposure [4, 5]. Destruction of limbal epithelial stem cells compromises corneal regeneration, resulting in persistent epithelial defects, conjunctivalization, neovascularization, stromal fibrosis, chronic inflammation, and progressive visual impairment [6, 7]. In parallel, dysfunction of the lacrimal gland, conjunctival goblet cells, and meibomian glands contributes to chronic dry eye disease.

Current clinical management of mustard keratopathy is largely supportive and focuses on suppressing inflammation with corticosteroids such as dexamethasone [8]. Although effective during the acute inflammatory phase, prolonged corticosteroid therapy is associated with significant complications, including elevated intraocular pressure, cataract formation [9], and delayed epithelial wound healing [10]. Despite considerable efforts to develop medical countermeasures, including anti-inflammatory, antioxidant, anti-fibrotic, and anti-angiogenic therapies, no treatment effectively targets the underlying mechanisms responsible for mustard-induced tissue injury or prevents long-term disease progression.

Accumulating evidence identifies mitochondrial dysfunction as a central driver of oxidative stress and epithelial injury following vesicant exposure. Mitochondria represent the major intracellular source of reactive oxygen species, and excessive mitochondrial reactive oxygen species production initiates bioenergetic failure, oxidative damage, and apoptotic cell death. Previous studies have demonstrated that mustard agents [11–14] directly damage mitochondria, yet the molecular pathways linking mitochondrial dysfunction to corneal pathology remain poorly understood, limiting the development of mechanism-based therapies.

Our laboratory previously identified mitochondrial dysfunction as a promising therapeutic target for ocular surface disease. We demonstrated that dynasore and dyngo-4a, small molecules that inhibit classic dynamin DNM2-mediated membrane remodeling, potently protect against tert-butyl hydrogen peroxide (tBHP)-induced injury in a human corneal epithelial cell model [15, 16]. Mechanistic studies revealed that dynasore suppresses calcium influx, prevents mitochondrial permeability transition pore opening, and promotes adaptive unfolded protein response signaling, thereby preserving mitochondrial function and epithelial survival [17–21]. Importantly, dynasore also reduced epithelial damage in a mouse model of dry eye disease, establishing modulation of mitochondrial stress pathways as a viable therapeutic strategy for ocular surface disorders.

Surprisingly, dynasore failed to protect corneal epithelial cells from nitrogen mustard-induced injury, despite its efficacy against other oxidative insults [19]. In contrast, mitochondrial division inhibitor-1 (Mdivi-1), a small molecule that specifically targets the dynamin DNM1L, produced robust cytoprotection. DNM1L (aka Drp1) is a mitochondria-localized GTPase that acts as the master regulator of mitochondrial fission, a process essential for mitochondrial quality control [22–25]. The protection of nitrogen mustard-exposed corneal epithelial cells by Mdivi-1 suggests that nitrogen mustard exposure injury is driven by mitochondrial dysfunction, and implicates mitochondrial fission as a critical therapeutic target.

In the present study, we comprehensively define mitochondrial mechanisms underlying nitrogen mustard-induced corneal injury in both our cell culture model in vitro and a mouse model in vivo. We demonstrate that nitrogen mustard rapidly induces mitochondrial fragmentation, bioenergetic collapse, membrane depolarization, oxidative stress, intracellular acidification, mitophagy, and apoptotic cell death. Pharmacologic inhibition of DNM1L with Mdivi-1 preserves mitochondrial structure and function, restores cellular metabolism, reduces oxidative damage, and markedly improves corneal epithelial integrity and tissue repair following nitrogen mustard exposure. Collectively, these findings establish mitochondrial dysfunction as a central pathological mechanism in mustard keratopathy and identify DNM1L-mediated mitochondrial remodeling as a therapeutically actionable target. Our work provides early preclinical evidence supporting mitochondrial-directed therapy as a promising strategy for treating mustard keratopathy.

## 2. Materials and methods

### 2.1. Chemical reagents

Nitrogen mustard (mechlorethamine HCl; Cat # 122564-5G) was purchased from Millipore Sigma (St. Louis, MO, USA). A stock solution was made (100 mM stock in DMSO), aliquoted, and stored at -70 degrees C. When needed for an experiment, a fresh aliquot was thawed and diluted into cell culture medium or with PBS to the appropriate concentration. The DMNL1 inhibitor Mdivi-1 (Cat # 475856-10MG), was purchased from Sigma-Aldrich (St. Louis, MO, USA). The more selective DMNL1 inhibitor, Drp1i27 dihydrochloride (Cat. No.: HY-152086A), was purchased from MedChem Express (Monmouth Junction, NJ, USA). These agents were also aliquoted as stock solutions, stored at -70 degrees C, and a fresh aliquot was used for all experiments.

Dexamethasone ophthalmic solution 0.1% (5 mL, generic) was procured from Bausch + Lomb (Bridgewater, NJ, USA). Fluorescein disodium salt (Cat# J61549.22) was purchased from ThermoFisher Scientific (Waltham, MA, USA). Hoechst 33342 trihydrochloride trihydrate dye (Cat# H1399) was obtained from Thermo Fisher Scientific.

### 2.2. Antibodies

Rabbit antibodies against TOM20, phosphorylated DNM1L (pSer616) and K12 were obtained from Thermo Fisher Scientific. Goat anti-rabbit IgG (H+L), highly cross-adsorbed secondary antibody conjugated to Alexa Fluor™ 647 (Cat# A-21245) was obtained from Thermo Fisher Scientific.

### 2.3. Human cell culture model of mustard keratopathy

For these studies, we used a cell culture model of mustard keratopathy that we previously described [19]. The model employs an immortalized human corneal limbal epithelial (HCLE) cell line [26], developed in the laboratory of Dr. Ilene Gipson, Harvard Medical School, according to methods described [27]. Cells were authenticated in the Gipson laboratory by marker expression analysis [28] and by chromosomal analysis using polymorphic short tandem repeat loci.

HCLE cells were cultured in Keratinocyte Serum-Free Medium (Cat# 17005042, ThermoFisher Scientific, Boston, MA, USA). For an experiment, the cells were seeded into a 12- or 24-well plate and maintained under standard culture conditions until they attained ∼70–80% confluency.

To create the mustard keratopathy model, nitrogen mustard stock solution was diluted into the culture medium of each plate well to a final concentration of 200 µM. Stock solutions of the treatment agents Mdivi-1 or Drp1i27 stock solutions were then diluted into the culture medium to the desired concentration. Cells were incubated for 2h. Following this time period, medium containing nitrogen mustard and treatments was removed and cells were washed with fresh Keratinocyte Serum-Free Medium.

For therapeutic outcome assessment by 4-[3-(4-iodophenyl)-2-(4-nitrophenyl)-2H-5-tetrazolio]-1,3-benzene disulfonate) assay (WST-1; Cat# MK400, Takara Bio, San Jose, CA, USA) assay, cells were allowed to recover for 18-20h. For the other endpoint assays described below, cells were assessed immediately following the 2h treatment period without an additional recovery period.

In the WST-1 assay, cellular metabolic activity is quantified by measurement of superoxide anions generated through NAD(P)H-dependent oxidoreductase activity. Anions reduce WST-1 to a soluble formazan product that exhibits absorbance in the visible range [29]. As WST-1 is cell-impermeable, this reduction predominantly occurs at the plasma membrane via extracellular electron transport. Formazan production was quantified by measuring absorbance at 450 nm using a BioTek Synergy H1 Microplate Reader (Winooski, VT, USA).

We previously established the optimal working concentration of mDivi-1, using a dose-response assay [19]. To do the same for Drpi27, HCLE cells exposed to 200 μM nitrogen mustard were treated with increasing concentrations of Drp1i27, and cell viability was assessed using the WST-1 assay. Drp1i27 improved cell viability in a concentration-dependent manner, with 50 μM providing the greatest protection against nitrogen mustard-induced cytotoxicity (Fig S1).

### 2.4. Immunostaining of cells

Indirect immunostaining of HCLE cells was performed using antibodies against TOM20 and pDNML1 Ser616. Cells were counterstained with DAPI to visualize nuclei. Imaging was performed using a Biotek Lionheart FX Automated Microscope (Agilent Technologies, Santa Clara, CA, USA).

### 2.5. MitoTracker and JC-1 staining

Mitochondrial morphology and membrane potential were assessed using MitoTracker™ Green FM (Thermo Fisher Scientific; Cat. No. M7514) and JC-1 (Thermo Fisher Scientific; Cat. No. T3168), respectively. HCLE cells were cultured in 24-well plates to sub-confluence and exposed to nitrogen mustard in the absence or presence of Mdivi-1 or Drp1i27.

To assess mitochondrial morphology, cells were incubated with MitoTracker in pre-warmed culture medium for 30 min at 37°C in the dark, washed with PBS, fixed with 4% paraformaldehyde for 10–15 min, and counterstained with Hoechst 33342 where indicated.

To assess mitochondrial membrane potential (ΔΨm), cells were incubated with JC-1 according to the manufacturer’s instructions, washed with PBS, and imaged immediately in fresh culture medium. Fluorescence images were acquired directly from 24-well plates using a Biotek Lionheart FX Automated Microscope under identical acquisition settings for all experimental groups. MitoTracker staining was used to evaluate mitochondrial network morphology and fragmentation, whereas the JC-1 red-to-green fluorescence intensity ratio was used as an indicator of mitochondrial membrane potential, with a reduced ratio reflecting mitochondrial depolarization.

### 2.6. MitoSOX staining for mitochondrial superoxide detection

Mitochondrial superoxide production was evaluated using the MitoSOX™ Red Mitochondrial Superoxide Indicator (Thermo Fisher Scientific, Waltham, MA, USA). Human corneal limbal epithelial (HCLE) cells were seeded in 24-well plates and cultured to approximately 70-80% confluence before exposure to nitrogen mustard in the presence or absence of Mdivi-1. Following treatment, cells were washed with pre-warmed Hank’s Balanced Salt Solution and incubated with 5 μM MitoSOX™ Red in Hank’s Balanced Salt Solution for 10 min at 37°C in the dark. Cells were then washed three times with HBSS to remove excess dye and immediately imaged using a Biotek Lionheart FX Automated Microscope under identical acquisition settings for all experimental groups. MitoSOX fluorescence intensity was quantified using NIH ImageJ from multiple randomly selected fields per well following background subtraction and normalized to the untreated control group.

### 2.7. Total antioxidant capacity assay

The intrinsic antioxidant capacity of Mdivi-1 was determined using the OxiSelect™ Total Antioxidant Capacity Assay Kit (Cell Biolabs, Inc., San Diego, CA, USA) according to the manufacturer’s protocol. Briefly, serial concentrations of Mdivi-1 were assayed alongside uric acid and glucose as positive and negative controls, respectively. Samples and standards were added to a 96-well microplate, followed by the assay reaction reagent. After incubation at room temperature, the reaction was terminated with stop solution, and absorbance was measured at 490 nm using a microplate reader. Total antioxidant capacity was calculated from a uric acid standard curve and expressed as uric acid equivalents. All experiments were performed in triplicate.

### 2.8. Mitophagy staining and assessment of mitochondrial biogenesis

Mitophagy was assessed using the Mitophagy Detection Kit (Dojindo Laboratories; Cat. No. MD01-10). HCLE cells were washed twice with Hanks’ HEPES buffer or serum-free medium and incubated with 100 nM Mtphagy Dye for 30 min at 37 °C. Following dye loading, cells were washed and exposed to nitrogen mustard in the presence or absence of Mdivi-1 or Drp1i27 for 2 h. To visualize lysosomes, cells were subsequently incubated with 1 μM Lyso Dye for 30 min at 37 °C, washed, and imaged directly in 24-well plates using a BioTek Lionheart FX imaging system under identical acquisition settings. Mitophagy was quantified by measuring Mtphagy Dye fluorescence and its co-localization with Lyso Dye, with increased red puncta and red-green co-localization indicating enhanced mitophagic activity.

Mitochondrial biogenesis was evaluated using the NovaQUANT® Mouse Mitochondrial to Nuclear DNA Ratio Kit (MilliporeSigma; Cat. No. 72621-M) according to the manufacturer’s instructions. Total genomic DNA was isolated from treated cells, and quantitative PCR was performed to determine mitochondrial DNA (mtDNA) and nuclear DNA (nDNA) content. The mtDNA ratio was calculated and used as an index of mitochondrial biogenesis across experimental groups.

### 2.9. pCAG-CAT-MitoTimer transfection

HCLE cells were seeded in 12-well plates and cultured to approximately 60% confluence before transfection with the pCAG-CAT-MitoTimer expression construct [30] using Lipofectamine™ 3000 Transfection Reagent (Thermo Fisher Scientific; Cat. No. L3000015) according to the manufacturer’s protocol. Briefly, plasmid DNA and Lipofectamine™ 3000 were diluted in Opti-MEM™ Reduced Serum Medium (Thermo Fisher Scientific; Cat. No. 51985091), mixed to allow formation of DNA-lipid complexes, and added to the cells.

Following transfection, cells were maintained under standard culture conditions to permit MitoTimer expression before exposure to nitrogen mustard in the presence or absence of Mdivi-1. Fluorescence images were acquired using a Biotek Lionheart FX Automated Microscope under identical acquisition settings for all experimental groups. MitoTimer was detected by the green (ex/em 488/518 nm) and red (ex/em 543/572 nm) channels. MitoTimer green fluorescence was used to identify newly synthesized mitochondria, whereas red fluorescence indicated aged or oxidized mitochondria. The green-to-red fluorescence intensity ratio was quantified as an index of mitochondrial turnover and oxidative status using ImageJ software.

### 2.10. pHrodo™ intracellular pH assay

Intracellular acidification was assessed using the pHrodo™ Red Intracellular pH Indicator AM dye (Thermo Fisher Scientific) to evaluate changes in cytosolic pH following nitrogen mustard exposure. HCLE cells were cultured in 24-well plates to approximately 70–80% confluence and treated with nitrogen mustard in the presence or absence of Mdivi-1. The pHrodo™ Red AM staining solution was prepared according to the manufacturer’s instructions by mixing the dye with PowerLoad™ concentrate and diluting the mixture in Live Cell Imaging Solution (LCIS; Thermo Fisher Scientific; Cat. No. A14291DJ). Cells were washed once with LCIS and incubated with the staining solution for 30 min at room temperature in the dark. Following staining, cells were washed with LCIS, maintained in fresh imaging buffer, and imaged immediately using a Biotek Lionheart FX Automated Microscope under identical acquisition settings for all experimental groups. Fluorescence intensity was quantified using NIH ImageJ following background subtraction. Because pHrodo™ Red fluorescence increases as intracellular pH decreases, increased fluorescence intensity was interpreted as enhanced intracellular acidification.

### 2.11. Gene expression quantification

Total RNA was isolated from HCLE cells treated with nitrogen mustard in the presence or absence of 50 μM Mdivi-1 using the RNeasy Plus Micro Kit (Qiagen; Cat. No. 74104) according to the manufacturer’s instructions. RNA concentration and quality were assessed using a Synergy H1 Microplate Reader (BioTek, Winooski, VT, USA). Complementary DNA (cDNA) was synthesized from purified RNA using the SMART-Seq® HT Kit (Takara Bio Inc.; Cat. No. 634456) following the manufacturer’s protocol.

Quantitative real-time PCR (qRT-PCR) was performed on a CFX Connect Real-Time PCR Detection System (Bio-Rad, Hercules, CA, USA) using PowerUp™ SYBR™ Green Master Mix (Thermo Fisher Scientific; Cat. No. 4309155). Gene expression analysis included transcripts involved in oxidative stress and mitochondrial regulation, including eNOS and catalase. The primer sequences used were:

### CAT (catalase)

- Forward: TGGGATCTCGTTGGAAATAACAC
- Reverse: TCAGGACGTAGGCTCCAGAAG

### NOS3 (nitric oxide synthetase 3)

- Forward: TGATGGCGAAGCGAGTGAAG
- Reverse: ACTCATCCATACACAGGACCC

Amplification specificity was verified by melt curve analysis. All reactions were performed in technical triplicate, and relative gene expression was calculated using the ΔΔCt method with GAPDH as the endogenous reference gene.

### 2.12. Seahorse assays

Mitochondrial bioenergetics were evaluated using the Seahorse XF Mito Stress Test on a Seahorse XF Pro Analyzer (Agilent Technologies, Santa Clara, CA, USA). HCLE cells were seeded in Seahorse XF cell culture microplates and cultured in Keratinocyte Serum-Free Medium (Thermo Fisher Scientific; Cat. No. 17005042) to approximately 80% confluence. Cells were treated with nitrogen mustard alone, nitrogen mustard in combination with Mdivi-1 (12.5, 25, or 50 μM), or 50 μM Mdivi-1 alone for 1 h. Vehicle controls containing equivalent concentrations of DMSO were included in all experiments.

The Seahorse XF sensor cartridge was hydrated overnight in Seahorse XF Calibrant (Agilent; Cat. No. 100840-000) at 37 °C in a non-CO₂ incubator according to the manufacturer’s instructions. On the day of the assay, culture medium was replaced with Seahorse XF assay medium (Agilent; Cat. No. 103680-100) supplemented with 10 mM glucose, 2 mM L-glutamine, and 1 mM sodium pyruvate (pH 7.4), and cells were equilibrated for 1 h at 37 °C in a non-CO₂ incubator.

Mitochondrial respiration was measured using the Seahorse XF Cell Mito Stress Test Kit (Agilent; Cat. No. 103015-100). Oxygen consumption rate (OCR) was recorded under basal conditions and following sequential injections of oligomycin (1.5 μM), FCCP (1.0 μM), and rotenone/antimycin A (0.5 μM). Basal respiration, ATP-linked respiration, maximal respiration, spare respiratory capacity, proton leak, and non-mitochondrial respiration were calculated using Wave software (Agilent Technologies). OCR values were normalized to cell number to account for differences in cell density among treatment groups.

### 2.13. Mouse model of mustard keratopathy

All animal experiments conformed to the ARVO Statement for the Use of Animals in Ophthalmic and Vision Research and to the recommendations of the National Institutes of Health Guide for the Care and Use of Laboratory Animals. The study was in compliance with ARRIVE guidelines. Breeding and animal procedures were approved by the IACUC of Tufts University. All mice were housed in a temperature- and humidity-controlled vivarium under a 12 h light/dark cycle with ad libitum access to food and water.

For initial experiments, we employed C57BL/6J mice (Strain code: 027), purchased from Charles River Laboratories (Wilmington, MA, USA).

For later experiments, we employed CAG-CAT-MitoTimer conditional transgenic mice on the C57Bl/6J background [31]. This strain carries the pMitoTimer reporter transgene, which encodes a mitochondrial-targeted redox-sensitive fluorescent protein that undergoes an irreversible shift from green fluorescence (GFP) to red fluorescence (DsRed) upon oxidation. The transgene is positioned downstream of a LoxP-flanked chloramphenicol acetyltransferase (CAT) STOP cassette under the control of the cytomegalovirus enhancer/chicken β-actin (CAG) promoter, allowing conditional expression following Cre-mediated recombination.

CAG-CAT-MitoTimer mice were crossed with tamoxifen-inducible CreERT2 mice purchased from Jackson Laboratories (Stock No. 004682) to generate double-transgenic offspring. MitoTimer⁺/Cre⁺ mice were used for all experiments. Expression of the MitoTimer reporter was induced by treating the mice with tamoxifen 10 days prior to an experiment (40 mg/kg, i.p.). Tamoxifen was injected intraperitoneally once daily for 10 days. This causes CreERT2 to translocate to the nucleus and excise the floxed STOP cassette, enabling expression of the mitochondrial MitoTimer reporter.

Genotyping was performed using genomic DNA isolated from mouse ear snips, according to the Tufts IACUC Policy on Identification Methods and Genotyping. Tail biopsies were performed to collect tissue for genotyping on mice younger than 17 days. Ethanol disinfected scissors were used to take a small snip of the tail for DNA. Following the procedure, the biopsy site was observed for active bleeding and hemostasis ensured before returning animals to the housing facilities. Primer sets previously described by Wilson et al. [30] were used to identify the MitoTimer and Cre transgenes. Genotyping analysis was conducted through the Transnetyx automated genotyping service (Transnetyx Inc., Cordova, TN, USA), and genotype assignments provided by the service were used to identify experimental animals.

In our first set of experiments, we used a previously described mouse model of mustard keratopathy [32]. In this model, a 2 μL drop of 40 mM nitrogen mustard in PBS was delivered to the central cornea as a topical drop and left for 2.5 min. This was followed by thorough PBS irrigation. Using this method, we found it difficult to control the spread of nitrogen mustard solution to surrounding tissues. Marked inflammation, eyelid closure, and in some cases severe ocular damage occurred. Outcome assessment using fluorescein staining and histology was possible, but the severity of injury handicapped detailed mechanistic analyses.

To limit nitrogen mustard exposure to the central cornea, we adopted a second delivery method. Rather than using a topical drop, nitrogen mustard solution (at various concentrations) was delivered to the central cornea by application of a 1.5 mm filter paper disc dampened with nitrogen mustard solution for 2.5 min. The disk was then removed and the cornea was not rinsed with PBS. After performing a titration study (Fig. S5), we selected 20 mM nitrogen mustard as the optimal concentration, causing clear tissue injury, while also enabling reliable downstream analyses of mitochondrial function and dysfunction.

Nitrogen mustard exposure procedures were conducted in anesthetized mice (ketamine (90-120 mg/kg) and xylazine (5-10 mg/kg), plus isoflurane inhalation (3-5% induction, 1-3% maintenance). To manage pain upon awakening of mice from anesthesia, buprenorphine ER LAB was administered subcutaneously to mice at 0.5 - 1.5 mg/kg volume based on body weight, one day prior to nitrogen mustard exposure and every 3 days after that for 2 weeks.

Following nitrogen mustard exposure, treatment, and clinical assessments, mice were returned to housing. Mice were euthanized at the end of each experiment by 100% CO2 inhalation followed by cervical dislocation. The CO2 flow rate for euthanasia systems was adjusted to displace 30% to 70% of the chamber volume/min. This method of euthanasia is consistent with recommendations of the Panel on Euthanasia of the American Veterinary Medical Association.

### 2.14. Preparation of Mdivi-1 eye drops and treatment of the ocular surface

Mdivi-1 eye drops were prepared by dissolving Mdivi-1 in DMSO to generate a 60 mM stock solution, which was stored at −20°C. Before use, the stock solution was diluted in sterile PBS to a final working concentration of 50 μM for topical administration. The final DMSO concentration in the eye drop formulation was maintained at ≤0.1%, a level that is well tolerated and not associated with corneal toxicity or irritation. DMSO 0.1% was used for vehicle treatment.

Two hours after nitrogen mustard exposure, the ocular surface was treated topically with Mdivi-1, vehicle, or dexamethasone ophthalmic solution 0.1% (Bausch + Lomb) eye drops, twice daily for 15 days.

### 2.15. Clinical assessment: fluorescein staining

Corneal epithelial damage and wound healing were evaluated in anesthetized mice by clinical fluorescein staining at days 1, 7, 14, 21, and 28 following nitrogen mustard exposure. Briefly, a 2 µL drop of 1% fluorescein disodium salt (prepared in PBS) was applied to the ocular surface and allowed to distribute for 1 min. Excess dye was gently rinsed away with sterile PBS before imaging. Clinical fluorescence images were acquired using the Phoenix Micron IV rodent imaging system (Phoenix Micron, Bend, OR, USA) with slit-lamp fluorescence settings under identical acquisition parameters for all eyes and time points. Fluorescein fluorescence intensity was quantified using NIH ImageJ software by measuring the integrated fluorescence within a standardized corneal region of interest following background subtraction.

### 2.16. Endpoint assessment: H&E

Following euthanasia, whole eyes were removed from mice and prepared for hematoxylin and eosin (H&E) staining. Eyes were fixed in 4% paraformaldehyde (PFA) in PBS at 4°C for 24 h, washed in PBS, dehydrated through a graded ethanol series, cleared in xylene, and embedded in paraffin. Paraffin blocks were sectioned at 5 µm using a rotary microtome and mounted onto glass slides.

Prior to H&E staining, sections were deparaffinized in xylene, rehydrated through graded ethanol to distilled water, stained with hematoxylin, differentiated and blued as needed, and counterstained with eosin. Sections were then dehydrated, cleared in xylene, and cover slipped using a permanent mounting medium. Histological images of the cornea were acquired using a CX33 microscope (Olympus, Shinjuku City, Tokyo, Japan) equipped with a QColor 5 digital camera (Olympus). All images were captured using identical microscope and camera settings to enable consistent histological comparisons among experimental groups.

### 2.17. Endpoint assessment: immunostaining and Mitotimer reporter imaging in tissue sections and whole mounts

Following euthanasia of mice, whole corneas were dissected and embedded for frozen sectioning or prepared for whole mount imaging. Indirect immunostaining of tissue sections was performed using an antibody against corneal keratin K12. Imaging was performed using a Biotek Lionheart FX Automated Microscope.

K12 immunostaining was detected using the cyan channel (ex/em 435/480 nm). MitoTimer was detected using the green (ex/em 488/518 nm) and red (ex/em 543/572 nm) channels. Identical acquisition parameters were used for every sample of the same tissue type. The green-to-red fluorescence intensity ratio was quantified as an index of mitochondrial turnover and oxidative status using ImageJ software by measuring the integrated fluorescence within a standardized corneal region of interest following background subtraction.

### 2.18. Statistical analysis

All in vitro experiments using mustard keratopathy cell culture model were performed with at least three independent biological replicates (n=3-4), and key findings were confirmed across two to three independent experimental runs to ensure reproducibility. Data are presented as mean ± standard error of the mean (SEM). Comparisons across multiple treatment groups (control, nitrogen mustard, nitrogen mustard +Mdivi-1, and Mdivi-1 alone) were analyzed using one-way ANOVA followed by appropriate post hoc testing in GraphPad Prism. A significance threshold of P < 0.05 was used for all analyses.

For in vivo studies, animals were randomly assigned to experimental groups, and outcome assessments were performed in a blinded manner to minimize bias. Sample sizes were determined based on prior experience with similar models, approximating an effect size (mean group differences divided by SD) of 2.5. Five eyes per group allow detection of effect sizes of 2.5, testing at 2-sided alpha=0.05, 80% power. Group comparisons were analyzed using ANOVA with P < 0.05 considered statistically significant.

### 2.19. Chemical safety

All experimental protocols involving nitrogen mustard were reviewed and approved by the Institutional Environmental Health & Safety (EH&S) office prior to initiation. Handling and administration of nitrogen mustard were conducted exclusively within a certified, externally vented chemical fume hood, and access to designated exposure areas was limited to trained and authorized personnel. Personnel adhered to strict safety procedures, including the use of double-layered nitrile gloves, lab coats, protective eyewear with side shields, and appropriate face coverings to minimize exposure risk. Nitrogen mustard was procured in minimal quantities, stored securely in accordance with manufacturer and institutional guidelines, and opened only under controlled hood conditions. All contaminated materials, including liquid and solid waste, were disposed of through a dedicated hazardous waste stream coordinated and managed by EH&S to ensure regulatory compliance and safe handling.

## 3. Results

### 3.1. Nitrogen mustard disrupts mitochondrial homeostasis and induces redox imbalance in corneal epithelial cells, while Mdivi-1 preserves mitochondrial function

#### 3.1.1. Nitrogen mustard-induced DNM1L-dependent mitochondrial fission is inhibited by Mdivi-1

To determine whether nitrogen mustard exposure induces mitochondrial fission in human corneal epithelial cells, HCLE cells were immunostained for the mitochondrial outer membrane protein TOM20 and phosphorylated DNM1L (pDNM1L Ser616), a marker of DNM1L activation. In untreated control cells, TOM20 staining revealed an elongated, interconnected mitochondrial network distributed throughout the cytoplasm, while pDNM1L Ser616 immunoreactivity was minimal or nearly absent (Fig. 1A–C′), indicating low basal mitochondrial fission activity. Following nitrogen mustard exposure, the mitochondrial network became highly fragmented, appearing as punctate and condensed TOM20-positive structures (Fig. 1D). This morphological change was accompanied by a marked increase in pDNM1L Ser616 staining (Fig. 1E), with prominent overlap between TOM20 and pDNM1L signals in merged images (Fig. 1F–F′), consistent with activation of pDNM1L -mediated mitochondrial fission. In contrast, treatment with the DNM1L inhibitor Mdivi-1 largely preserved the elongated mitochondrial morphology and substantially reduced pDNM1L Ser616 immunoreactivity (Fig. 1G–I′), indicating suppression of nitrogen mustard-induced DNM1L activation and mitochondrial fragmentation.

**Figure 1.**
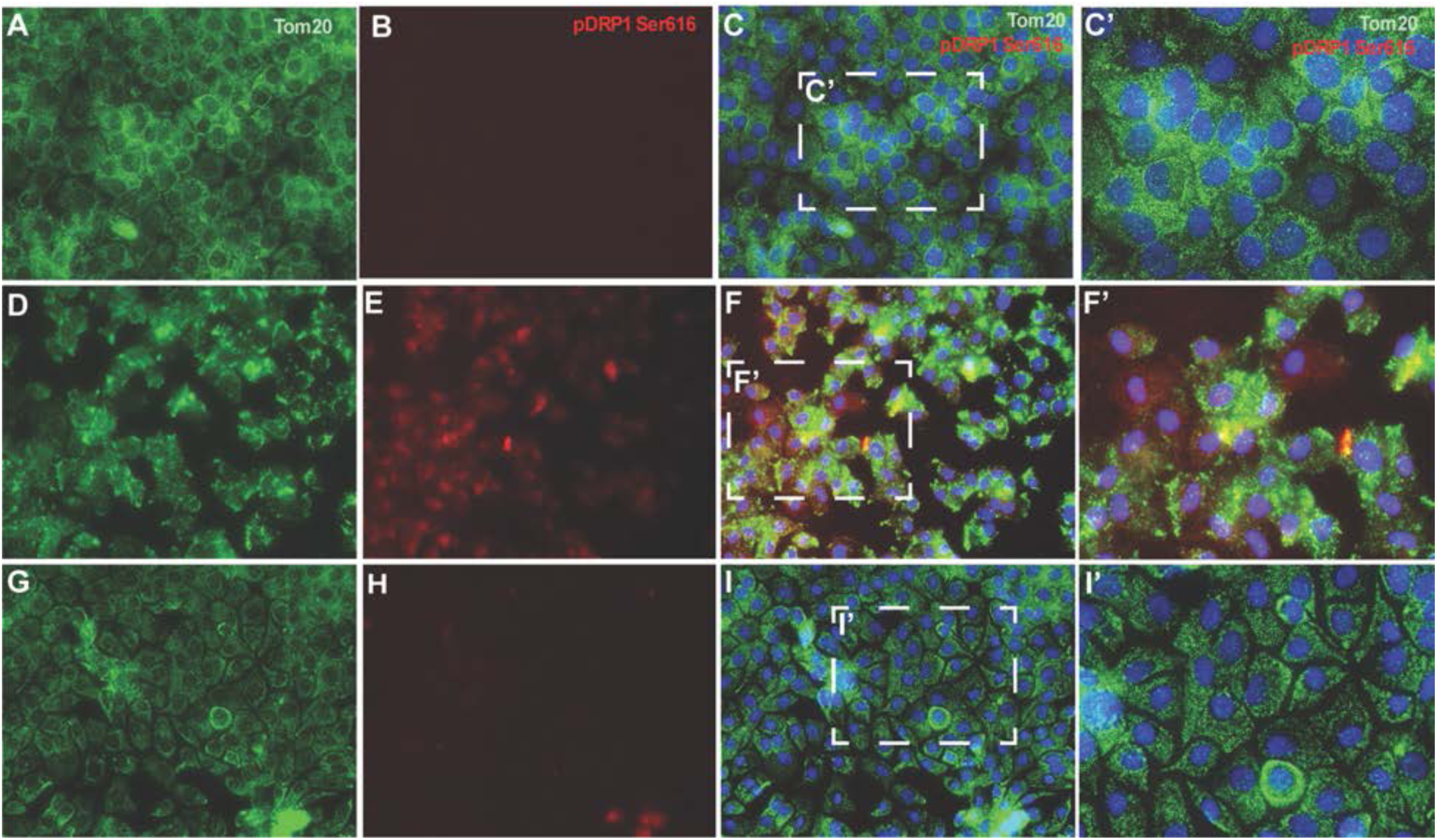
Nitrogen mustard activates DNM1L-dependent mitochondrial fission in corneal epithelial cells. Representative confocal images of HCLE cells immunostained for TOM20 (green), pDNML1 Ser616 (red), and DAPI (blue). (A-C′) Untreated control cells exhibit an elongated mitochondrial network with minimal pDNML1 Ser616 staining. (D-F′) Exposure to 200 µM nitrogen mustard induces mitochondrial fragmentation and markedly increases pDNML1 Ser616 expression. (G-I′) Mdivi-1 treatment preserves mitochondrial morphology and suppresses nitrogen mustard-induced pDNML1 Ser616 expression. C′, F′, and I′ are higher-magnification images of the boxed regions in C, F, and I, respectively.

#### 3.1.2. Mdivi-1 prevents nitrogen mustard-induced mitochondrial fragmentation and membrane depolarization

To determine whether Mdivi-1 protects against nitrogen mustard-induced alterations in mitochondrial morphology, HCLE cells were stained with MitoTracker Green and examined by fluorescence microscopy (Fig. 2). Untreated HCLE cells exhibited a well-organized mitochondrial network characterized by elongated and interconnected mitochondria distributed throughout the cytoplasm (Fig. 2A-C′). In contrast, exposure to 200 μM nitrogen mustard for 2 h caused marked mitochondrial fragmentation, characterized by numerous short, punctate mitochondria and disruption of the interconnected mitochondrial network (Fig. 2D-F′). Treatment with 50 μM Mdivi-1 substantially preserved mitochondrial morphology, maintaining a more elongated and interconnected mitochondrial network while reducing mitochondrial fragmentation compared with nitrogen mustard -exposed cells (Fig. 2G–I′). A similar result was observed using the more selective inhibitor of DNM1L, Drp1i27 (Fig. S2)

**Figure 2.**
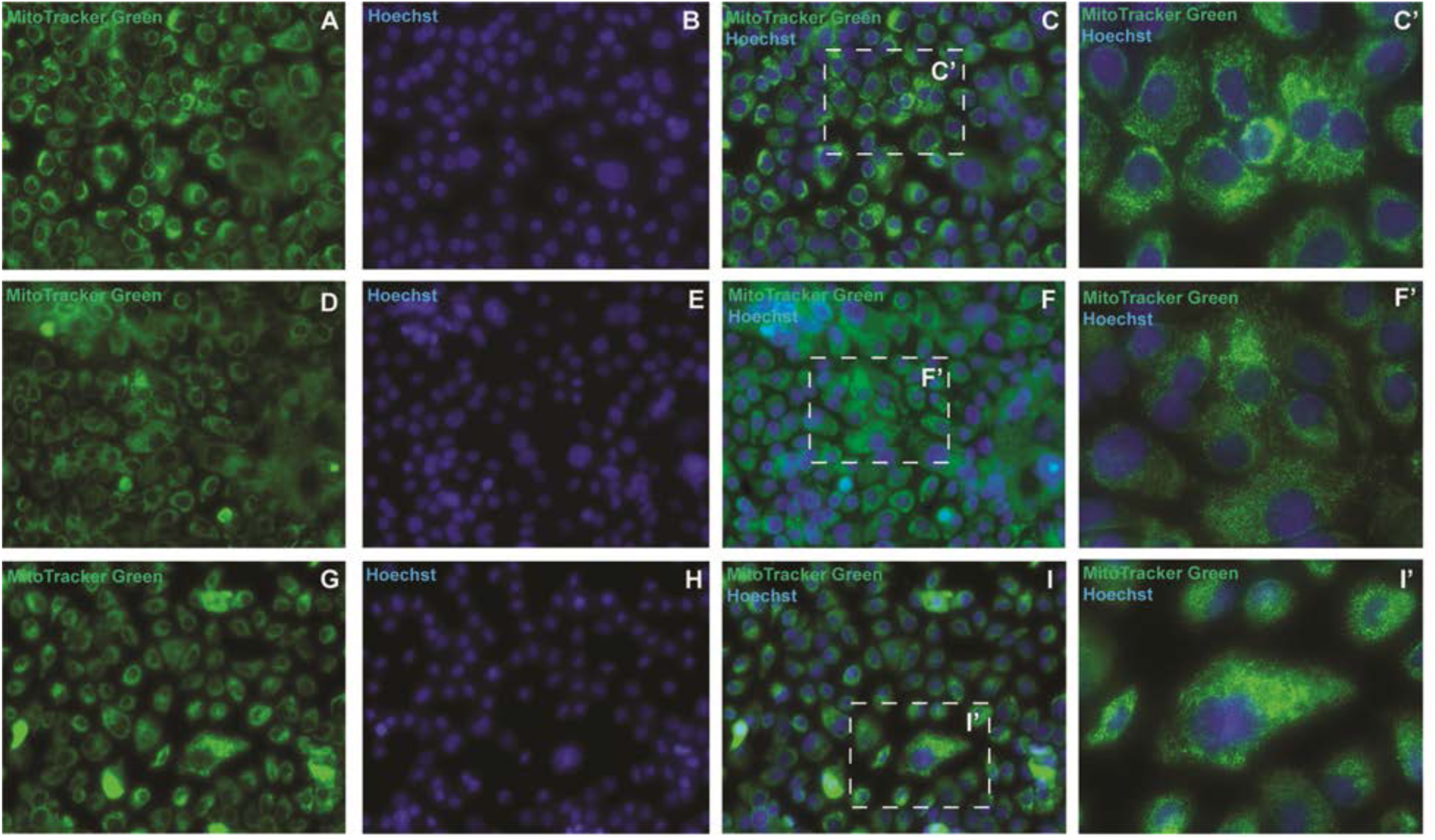
Mdivi-1 attenuates nitrogen mustard-induced mitochondrial fragmentation in HCLE cells. HCLE cells were exposed to 200 μM nitrogen mustard for 2 h and subsequently treated with 50 μM Mdivi-1. Mitochondrial morphology was assessed by MitoTracker staining. (A-C′) Untreated control cells showing an elongated and interconnected mitochondrial network. (D-F′) nitrogen mustard-treated cells exhibiting marked mitochondrial fragmentation and loss of network integrity. (G-I′) Mdivi-1 treatment attenuated nitrogen mustard-induced mitochondrial fragmentation and preserved mitochondrial morphology. Panels C′, F′, and I′ represent higher-magnification views of the boxed regions in C, F, and I, respectively.

Because mitochondrial morphology is closely linked to mitochondrial function, we next evaluated mitochondrial membrane potential (ΔΨm) using the JC-1 assay (Fig. 3). Untreated HCLE cells displayed strong red JC-1 aggregate fluorescence and relatively low green JC-1 monomer fluorescence, indicative of polarized and functionally intact mitochondria (Fig. 3A–C). In contrast, nitrogen mustard exposure markedly decreased red fluorescence and increased green fluorescence, indicating mitochondrial membrane depolarization (Fig. 3D–F). Mdivi-1 treatment partially restored JC-1 aggregate fluorescence while reducing JC-1 monomer fluorescence (Fig. 3G–I), consistent with preservation of mitochondrial membrane potential. Quantitative analysis of the JC-1 red/green fluorescence ratio confirmed a significant reduction following nitrogen mustard exposure, whereas Mdivi-1 treatment significantly restored the red/green ratio compared with nitrogen mustard-exposed cells (Fig. 3J). A similar result was observed using the more selective DNM1L inhibitor, Drp1i27 (Fig. S3).

**Figure 3.**
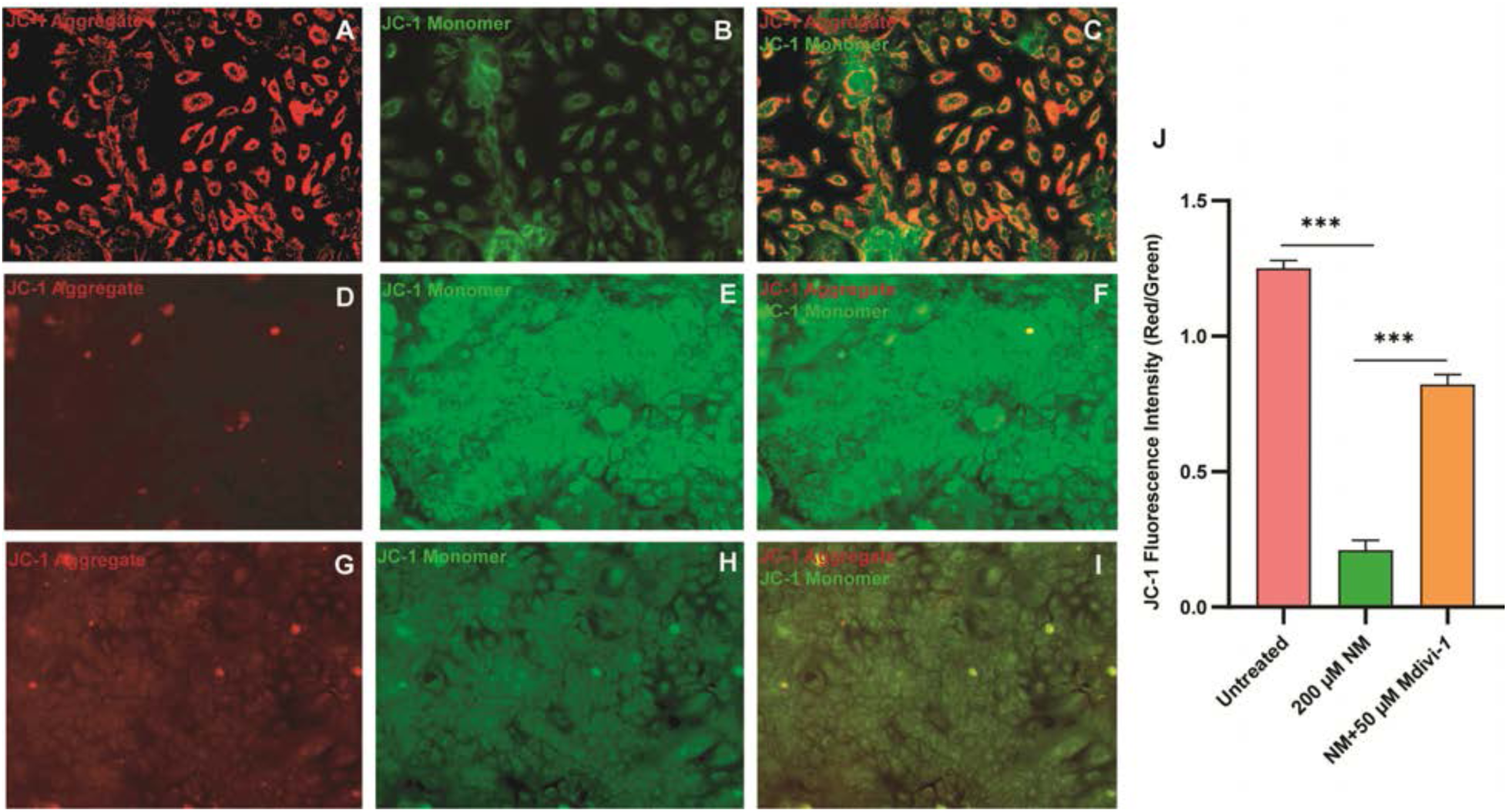
Mdivi-1 restores mitochondrial membrane potential following nitrogen mustard exposure in HCLE cells. Mitochondrial membrane potential (ΔΨm) was assessed using the JC-1 assay. Polarized mitochondria exhibit red JC-1 aggregates, whereas depolarized mitochondria display green JC-1 monomers. (A-C) Untreated control cells showing robust red fluorescence indicative of intact ΔΨm. (D-F) Nitrogen mustard-treated cells displaying increased green fluorescence and reduced red fluorescence, consistent with mitochondrial membrane depolarization. (G-I) Mdivi-1 treatment restored ΔΨm, evidenced by increased red fluorescence and reduced green fluorescence. (J) Quantification of the JC-1 red-to-green fluorescence intensity ratio demonstrating significant membrane depolarization following nitrogen mustard exposure and significant recovery following Mdivi-1 treatment. Data are presented as mean ± SEM. ***P < 0.001.

Together, these findings demonstrate that Mdivi-1 prevents nitrogen mustard-induced mitochondrial fragmentation and membrane depolarization in corneal epithelial cells and are consistent with a mechanism-of-action involving targeting of DNML1.

#### 3.1.3. Mdivi-1 preserves mitochondrial homeostasis by reducing nitrogen mustard-induced reactive oxygen species accumulation, antioxidant impairment, and mitochondrial permeability transition pore opening

Given that Mdivi-1 significantly reduced nitrogen mustard-induced mitochondrial reactive oxygen species accumulation, we next investigated the effect of Mdivi-1 on oxidative stress. MitoSOX staining revealed low basal mitochondrial superoxide levels in untreated HCLE cells (Fig. 4A–C). Exposure to 200 μM nitrogen mustard markedly increased MitoSOX fluorescence intensity (Fig. 4D–F), indicating substantial mitochondrial superoxide production. In contrast, treatment with 50 μM Mdivi-1 markedly reduced MitoSOX fluorescence (Fig. 4G–I), and quantitative analysis confirmed a significant decrease in mitochondrial superoxide levels compared with nitrogen mustard-exposed cells (Fig. 4J). A similar result was observed using the more selective inhibitor of DNM1L, Drp1i27 (Fig. S4).

**Figure 4.**
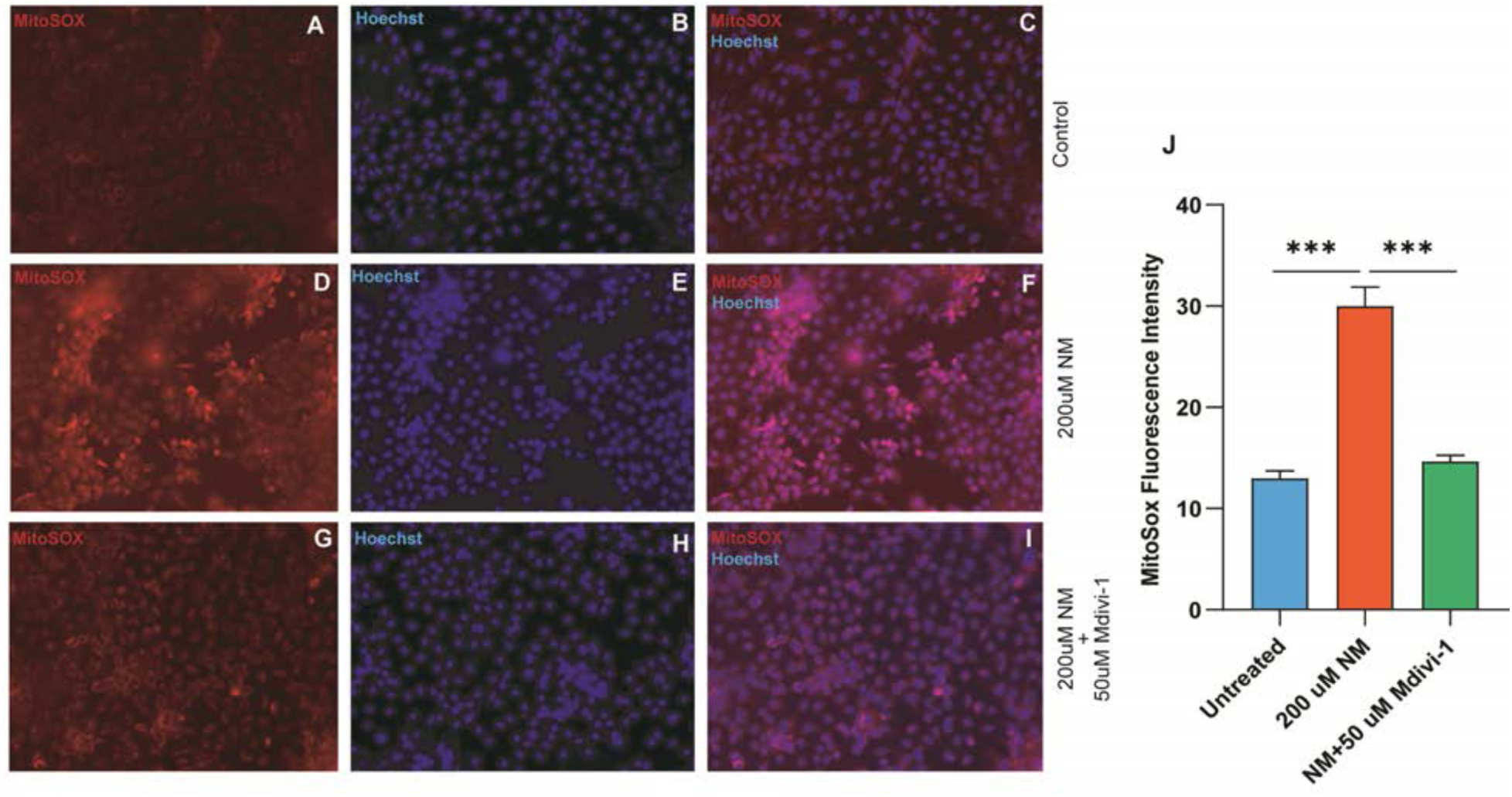
Mdivi-1 attenuates nitrogen mustard-induced mitochondrial ROS production in HCLE cells. Representative MitoSOX (red) and Hoechst (blue) fluorescence images of untreated cells (A-C), 200 μM NM-treated cells (D-F), and nitrogen mustard-exposed cells treated with 50 μM Mdivi-1 (G-I). Merged images are shown in C, F, and I. NM markedly increased MitoSOX fluorescence, indicating elevated mitochondrial superoxide production, whereas Mdivi-1 reduced mitochondrial ROS accumulation. (J) Quantification of MitoSOX fluorescence intensity. Data are presented as mean ± SEM. **\***P < 0.001.

To determine whether the reduction in mitochondrial reactive oxygen species was associated with changes in endogenous antioxidant defenses, we next examined the expression of antioxidant genes. Nitrogen mustard exposure significantly decreased the expression of catalase and eNOS compared with untreated cells (Fig. S5A, B). In contrast, Mdivi-1 treatment significantly restored the expression of both genes, increasing catalase and eNOS expression above the levels observed in nitrogen mustard-exposed cells (Fig. S5A, B).

Because Mdivi-1 reduced mitochondrial reactive oxygen species and restored antioxidant gene expression, we next asked whether these effects resulted from intrinsic antioxidant activity. A cell-free antioxidant assay was performed using uric acid as a positive control and glucose as a negative control. As expected, positive control uric acid exhibited strong concentration-dependent antioxidant activity, whereas negative control glucose showed negligible activity (Fig. S6). In comparison, Mdivi-1 displayed only minimal antioxidant activity over the tested concentration range, indicating that it is not an effective direct free radical scavenger.

To determine whether nitrogen mustard-induced oxidative stress progressed to irreversible mitochondrial injury, we next assessed mitochondrial permeability transition pore opening. Untreated HCLE cells exhibited strong mitochondrial calcein fluorescence, consistent with intact mitochondrial membrane integrity (Fig. 5A–C′). In contrast, nitrogen mustard exposure markedly reduced mitochondrial calcein fluorescence, indicating increased mitochondrial permeability transition pore opening (Fig. 5D–F′). Treatment with 50 μM Mdivi-1 substantially preserved mitochondrial calcein fluorescence and reduced mitochondrial permeability transition pore opening compared with nitrogen mustard-exposed cells (Fig. 5G–I′).

**Figure 5.**
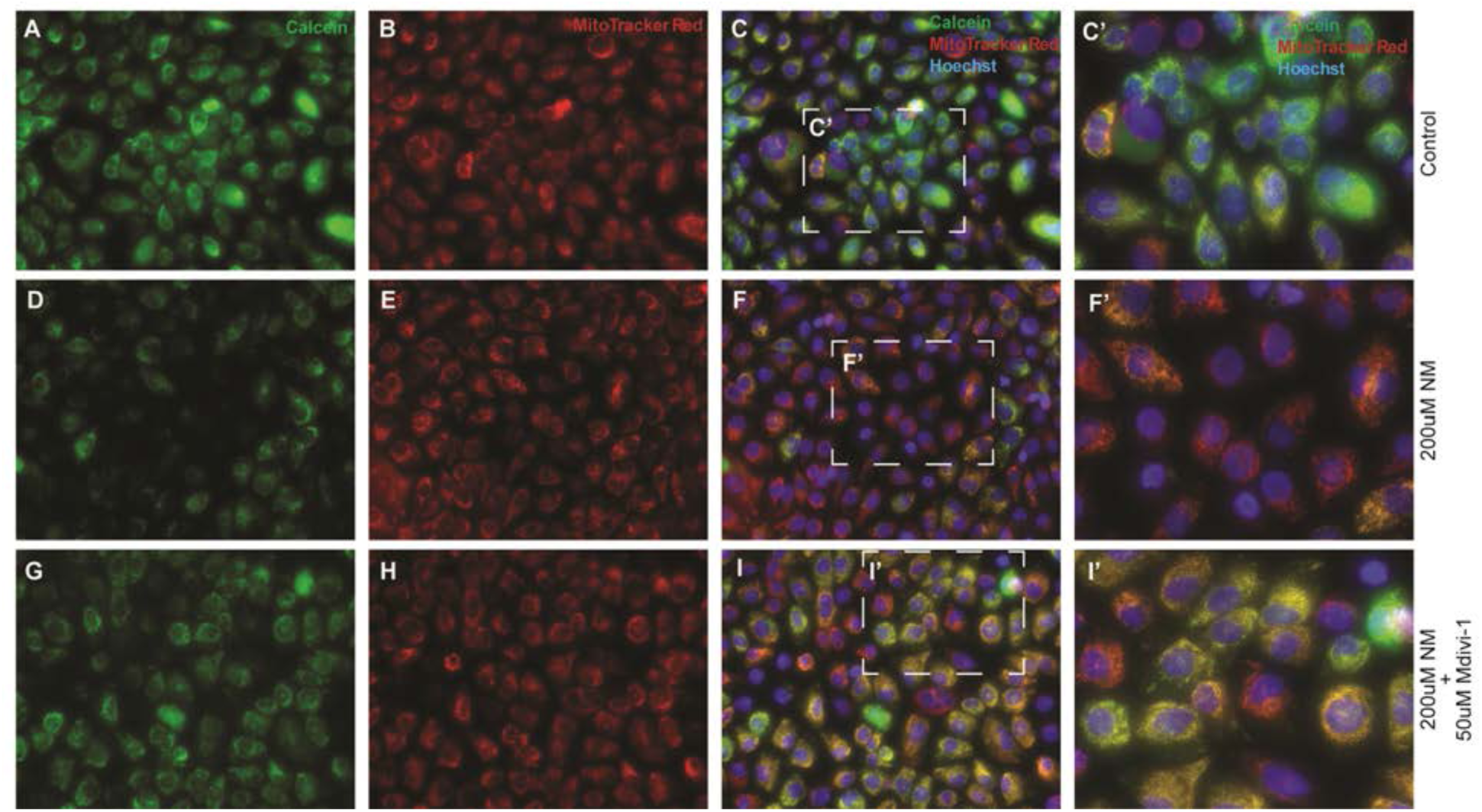
Mdivi-1 prevents nitrogen-induced mitochondrial permeability transition pore opening. Representative fluorescence images of HCLE cells stained with calcein (green), MitoTracker Red (red), and Hoechst (blue). Control cells (A-C), cells exposed to 200 µM nitrogen mustard (D-F), and cells treated with 200 µM nitrogen mustard plus 50 µM Mdivi-1 (G-I) are shown. Nitrogen mustard reduced mitochondrial calcein retention, indicating mitochondrial permeability transition pore opening, whereas Mdivi-1 restored mitochondrial calcein fluorescence. Boxed regions are enlarged in C′, F′, and I′.

Collectively, these results demonstrate that nitrogen mustard induces progressive mitochondrial dysfunction characterized by increased mitochondrial reactive oxygen species production, impaired endogenous antioxidant defenses, and increased mitochondrial permeability transition pore opening, whereas Mdivi-1 preserves mitochondrial integrity despite exhibiting only minimal intrinsic antioxidant activity.

### 3.2 Mdivi-1 restores nitrogen mustard-induced mitochondrial quality control and bioenergetic dysfunction

#### 3.2.1 Mdivi-1 enhances mitochondrial quality control in nitrogen mustard exposed HCLE cells by promoting mitophagy, mitochondrial biogenesis, and mitochondrial turnover

Damaged mitochondria that escape quality-control mechanisms become persistent sources of reactive oxygen species and bioenergetic dysfunction. Under physiological conditions, mitochondrial homeostasis is maintained through coordinated mitophagy and mitochondrial biogenesis, which together eliminate dysfunctional mitochondria and replenish the mitochondrial pool with newly synthesized organelles. We therefore next investigated whether the restoration of mitochondrial function by Mdivi-1 was accompanied by improvements in mitochondrial quality-control pathways.

Mitophagy was evaluated by assessing the colocalization of mitophagy-positive mitochondria with lysosomes, an indicator of mitolysosome formation (Fig. 6). nitrogen mustard-exposed HCLE cells exhibited limited colocalization between mitophagy and lysosomal signals, indicating reduced mitophagic activity (Fig. 6A–F′). In contrast, treatment with 50 μM Mdivi-1 markedly increased the colocalization of mitophagy-positive mitochondria with lysosomes, resulting in numerous yellow/orange puncta indicative of mitolysosome formation (Fig. 6G–J′). Higher-magnification images further demonstrated increased phagolysosome formation in Mdivi-1-treated cells (Fig. 6J′, arrows), consistent with enhanced mitophagy.

**Figure 6.**
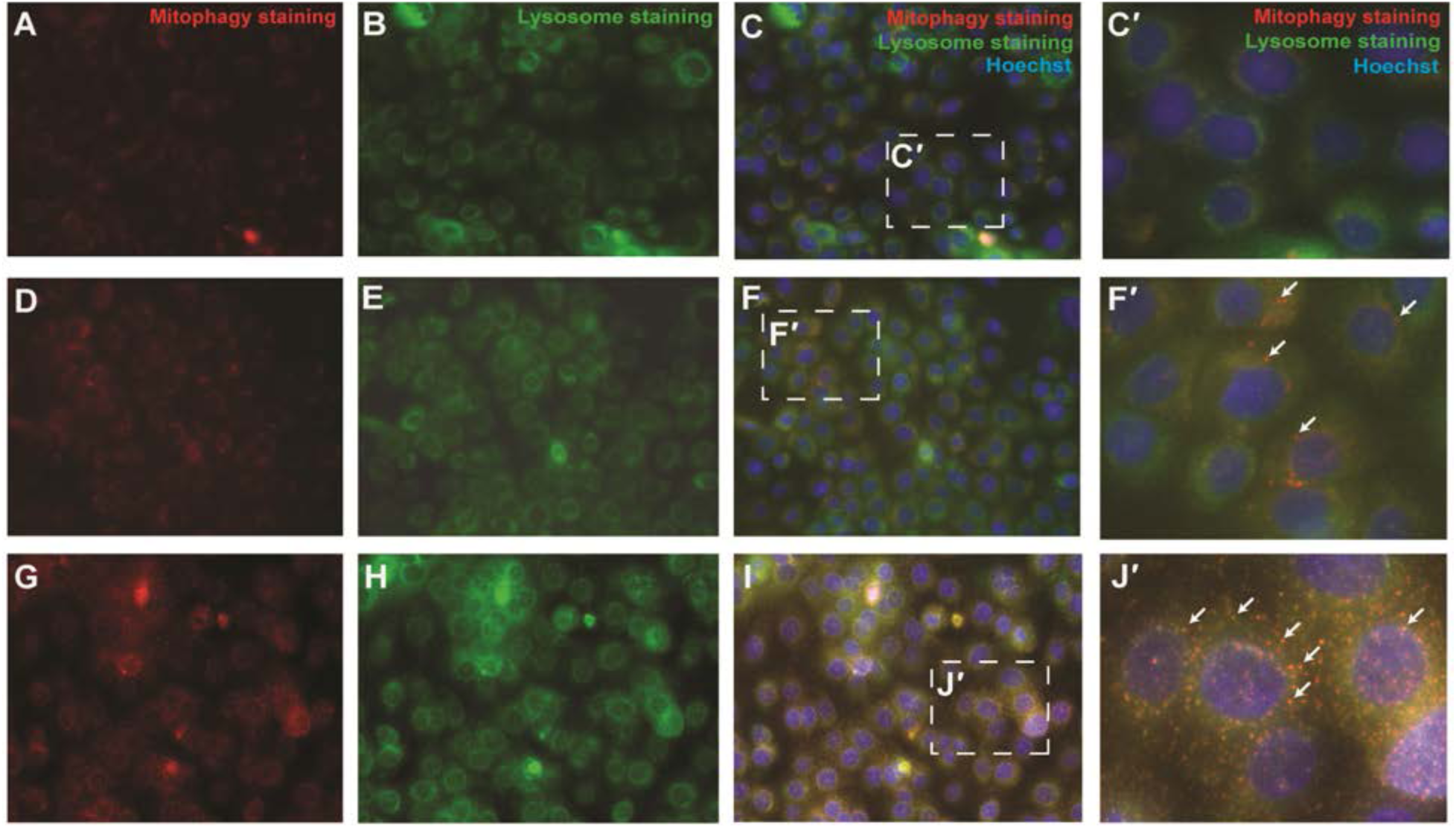
Mdivi-1 enhances mitophagy and phagolysosome formation following nitrogen mustard-induced injury in HCLE cells. Mitophagy (red), lysosomes (green), and nuclei (Hoechst, blue) were visualized by fluorescence microscopy. (A-C′) NM-treated cells. (D-F′) nitrogen mustard + 50 μM Mdivi-1-treated cells. Mdivi-1 increased the number of colocalized red and green puncta (yellow/orange), representing mitochondria-containing phagolysosomes (mitolysosomes) and indicating enhanced delivery of damaged mitochondria to lysosomal degradative pathways. Arrows highlight representative phagolysosomal structures. C′ and F′ represent higher-magnification views of the boxed regions.

Because efficient mitochondrial quality control requires replacement of eliminated mitochondria, we next assessed mitochondrial biogenesis by quantifying mitochondrial DNA copy number. Mdivi-1 treatment significantly increased mitochondrial DNA copy number compared with nitrogen mustard-exposed cells (Fig. S7), consistent with mitochondrial biogenesis.

We next assessed mitochondrial turnover using the MitoTimer reporter assay (Fig 7). Untreated HCLE cells exhibited predominantly green fluorescence, representing healthy mitochondria with minimal oxidative damage (Fig. 7A–C′). In contrast, nitrogen mustard exposure markedly increased red fluorescence and reduced green fluorescence, indicating the accumulation of oxidized and aged mitochondria (Fig. 7D–F′). Treatment with Mdivi-1 increased the proportion of green mitochondria while reducing red fluorescence (Fig. 7G–I′). Quantitative analysis confirmed that nitrogen mustard exposure significantly decreased the MitoTimer green-to-red fluorescence intensity ratio, whereas Mdivi-1 significantly restored this ratio (Fig. 7J), consistent with enhanced mitochondrial turnover.

**Figure 7.**
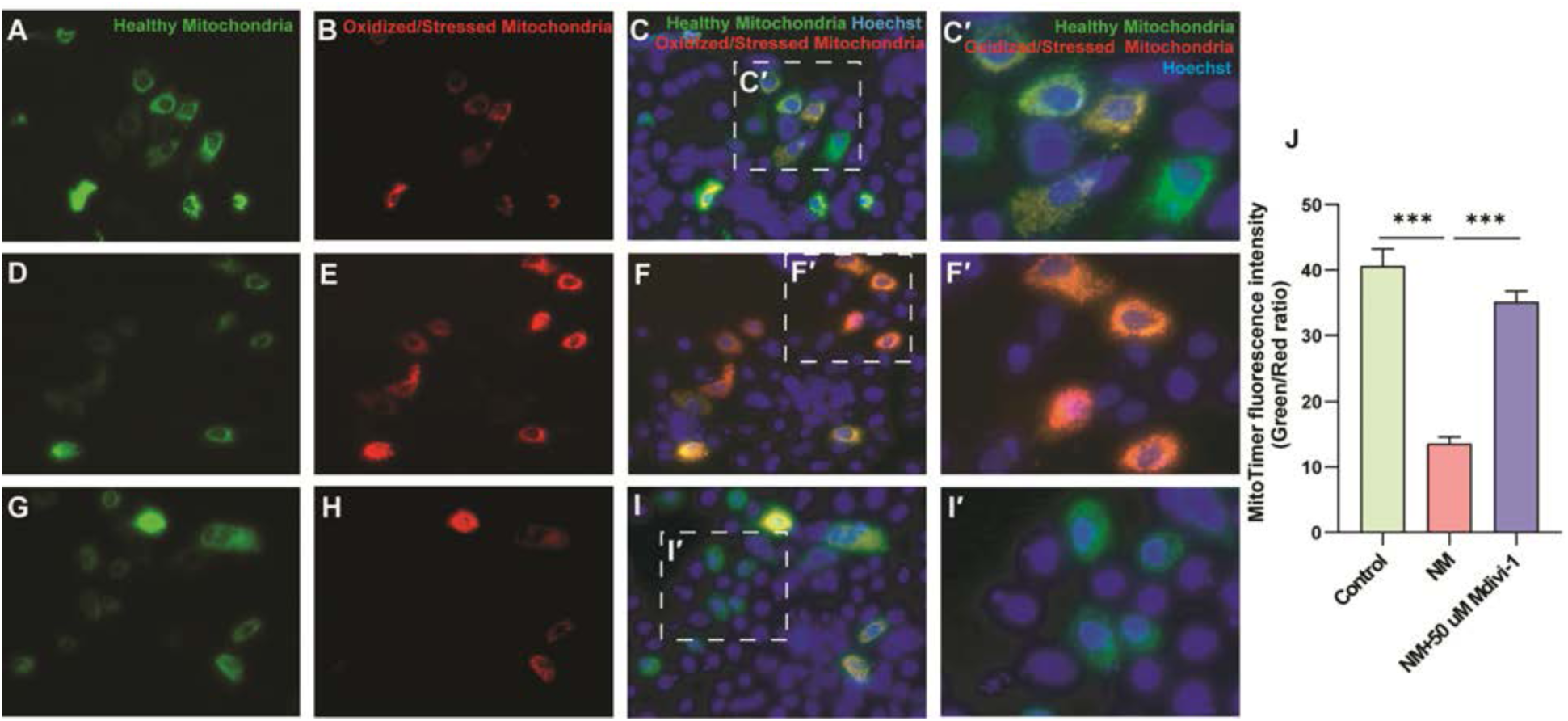
Mdivi-1 enhances mitochondrial turnover in MitoTimer-transfected HCLE cells. HCLE cells were transiently transfected with the MitoTimer plasmid to assess mitochondrial turnover. MitoTimer shifts from green to red fluorescence as mitochondria age and accumulate oxidative damage. **(A–C′)** Control cells. **(D–F′)** Nitrogen mustard-treated cells showing increased red fluorescence and accumulation of aged/oxidatively stressed mitochondria. **(G–I′)** nitrogen mustard + 50 μM Mdivi-1-treated cells displaying reduced red fluorescence and increased healthy mitochondrial populations, indicative of enhanced mitochondrial turnover and renewal. **(J)** Quantification of the MitoTimer green-to-red fluorescence intensity ratio. Nitrogen mustard significantly reduced the ratio, whereas Mdivi-1 significantly restored it, consistent with improved mitochondrial turnover. **C′, F′, and I′** represent higher-magnification views of the boxed regions. Data are presented as mean ± SEM. ***P < 0.001.

#### 3.2.2 Mdivi-1 restores mitochondrial respiration and prevents intracellular acidification

Having established that Mdivi-1 enhances mitochondrial quality control through mitophagy, mitochondrial biogenesis, and mitochondrial turnover, we next investigated whether these improvements translated into recovery of mitochondrial bioenergetic function. Mitochondrial respiration was assessed using the Seahorse Mitochondrial Stress Test following nitrogen mustard exposure and treatment with increasing concentrations of Mdivi-1 (Fig. 8A, B). Nitrogen mustard exposure markedly impaired mitochondrial respiration, as evidenced by significant reductions in basal respiration, maximal respiration, spare respiratory capacity, proton leak, and non-mitochondrial oxygen consumption compared with untreated controls (Fig. 8C–G). Treatment with Mdivi-1 improved all respiratory parameters in a dose-dependent manner. Compared with nitrogen mustard-exposed cells, Mdivi-1 significantly increased basal respiration, maximal respiration, spare respiratory capacity, proton leak, and non-mitochondrial oxygen consumption, with the greatest recovery observed at 50 μM (Fig. 8C–G).

**Figure 8.**
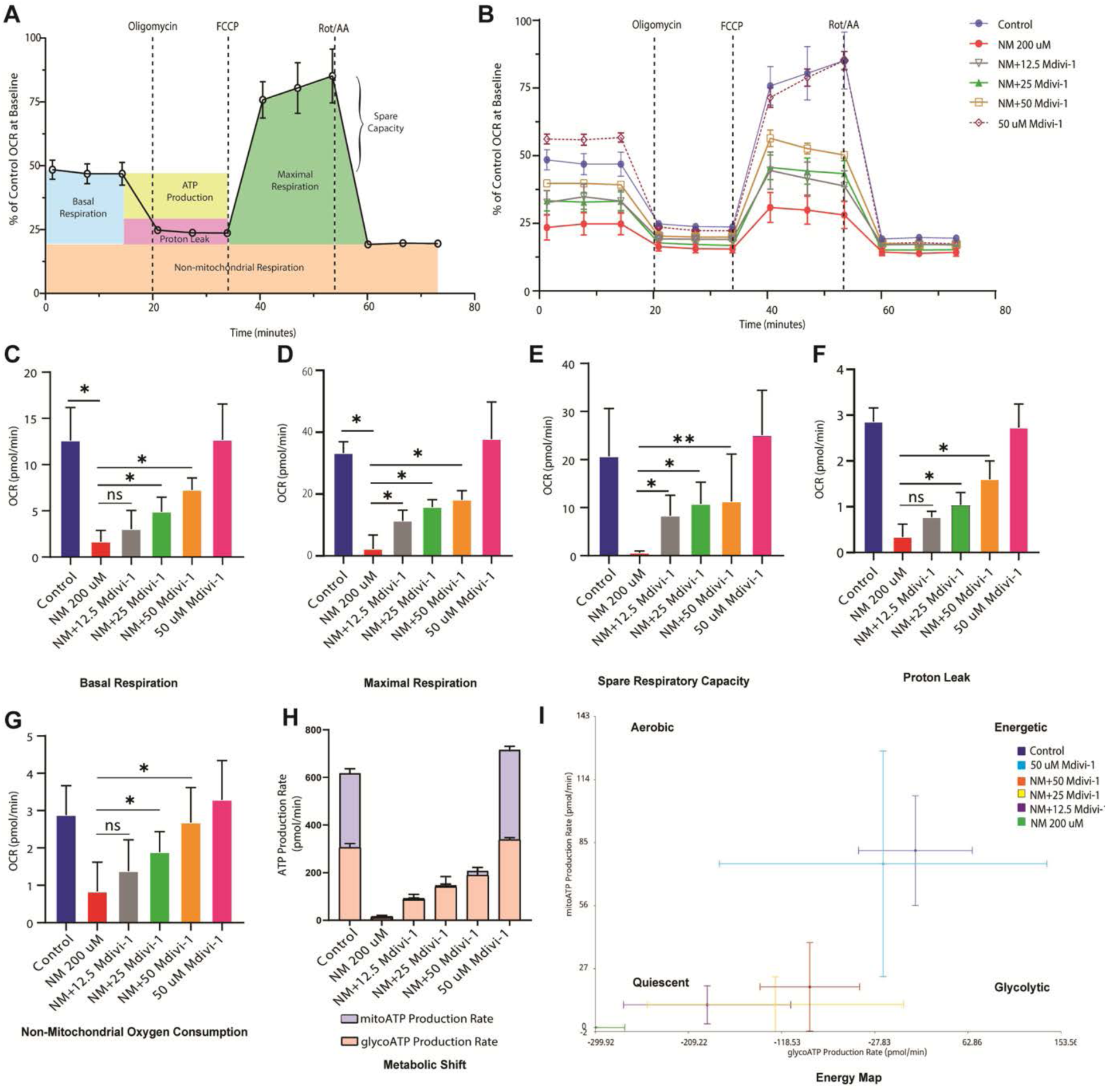
Mdivi-1 restores mitochondrial bioenergetic function following nitrogen mustard-induced injury in HCLE cells. (A) Schematic of the Seahorse XF mitochondrial stress test illustrating determination of basal respiration, ATP-linked respiration, proton leak, maximal respiration, spare respiratory capacity, and non-mitochondrial respiration following sequential addition of oligomycin, FCCP, and rotenone/antimycin A. (B) Representative oxygen consumption rate (OCR) profiles of control, nitrogen mustard-treated, and Mdivi-1-treated HCLE cells. (C–G) Quantification of basal respiration, maximal respiration, spare respiratory capacity, proton leak, and non-mitochondrial oxygen consumption, respectively. Nitrogen mustard significantly impaired mitochondrial respiratory function, whereas Mdivi-1 restored respiratory parameters in a dose-dependent manner. (H) Seahorse ATP-rate assay showing mitochondrial ATP (mitoATP) and glycolytic ATP (glycoATP) production rates. Under basal conditions, HCLE cells generated ATP through a balanced contribution of oxidative phosphorylation and glycolysis. Nitrogen mustard reduced mitochondrial ATP production and shifted cellular metabolism toward glycolysis, whereas Mdivi-1 partially restored mitochondrial ATP generation and promoted a metabolic shift toward oxidative phosphorylation. (I) Seahorse energy phenotype map demonstrating restoration of cellular energetic status following Mdivi-1 treatment. Data are presented as mean ± SEM. *P < 0.05, **P < 0.01.

To further evaluate the effects of Mdivi-1 on cellular energy metabolism, ATP production rates were determined using the Seahorse ATP Rate Assay (Fig. 8H). Nitrogen mustard exposure markedly reduced mitochondrial ATP production, with glycolysis becoming the predominant source of cellular ATP. Mdivi-1 treatment dose-dependently increased mitochondrial ATP production while maintaining glycolytic ATP production. At 50 μM, Mdivi-1 partially shifted cellular energy metabolism toward mitochondrial ATP production, although glycolysis remained a substantial contributor to total ATP generation (Fig. 8H).

Consistent with these findings, Seahorse energy phenotype analysis demonstrated that nitrogen mustard exposure shifted HCLE cells toward a metabolically compromised, less energetic phenotype (Fig. 11I). In contrast, Mdivi-1 progressively shifted the metabolic phenotype toward a more energetic and aerobic state, approaching that of untreated control cells (Fig. 8I).

Because impaired mitochondrial respiration is frequently associated with intracellular acidification, we next assessed intracellular pH using the pH-sensitive fluorescent probe pHrodo (Fig. 9). Untreated HCLE cells exhibited low pHrodo fluorescence intensity (Fig. 9A–C), whereas nitrogen mustard exposure markedly increased pHrodo fluorescence, indicating intracellular acidification (Fig. 9D–F). Treatment with 50 μM Mdivi-1 reduced pHrodo fluorescence compared with nitrogen mustard -exposed cells (Fig. 9G–I). Quantitative analysis confirmed a significant increase in pHrodo fluorescence following nitrogen mustard exposure and a significant reduction following Mdivi-1 treatment (Fig. 9J).

**Figure 9.**
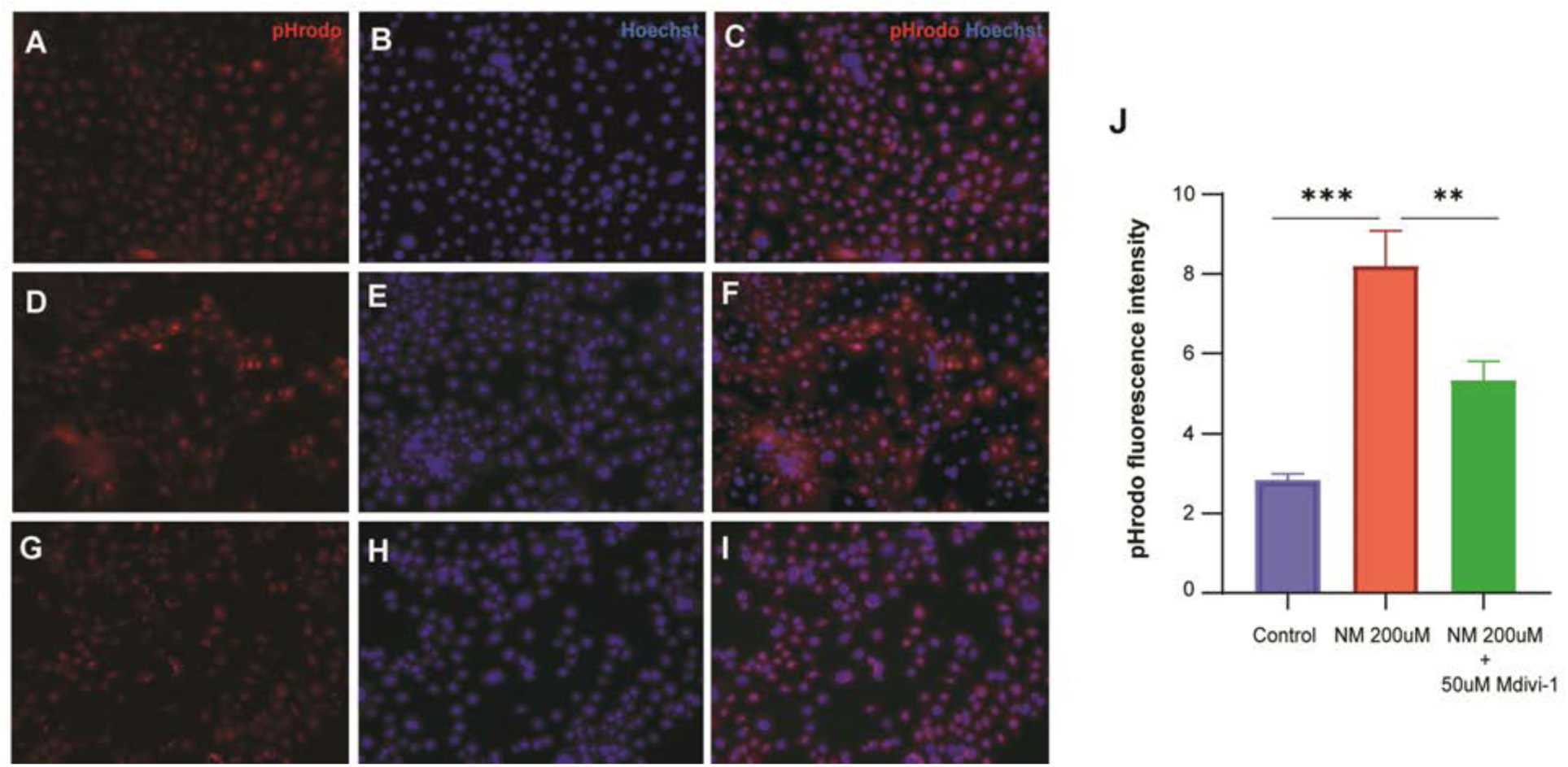
Mdivi-1 reduces intracellular acidification following nitrogen mustard -induced injury in HCLE cells. Intracellular acidification was assessed using the pH-sensitive fluorescent probe pHrodo (red); nuclei were counterstained with Hoechst (blue). (A–C) Untreated control cells exhibiting low pHrodo fluorescence. (D–F) nitrogen mustard -treated cells showing increased pHrodo fluorescence, indicative of intracellular acidification. (G–I) nitrogen mustard + 50 μM Mdivi-1-treated cells displaying reduced pHrodo fluorescence compared with nitrogen mustard -treated cells. (J) Quantification of pHrodo fluorescence intensity demonstrating significantly increased intracellular acidification following nitrogen mustard exposure and significant attenuation by Mdivi-1 treatment. Data are presented as mean ± SEM. **P < 0.01, ***P < 0.001.

Collectively, these findings demonstrate that Mdivi-1 restores mitochondrial bioenergetic function and alleviates nitrogen mustard-induced metabolic dysfunction in corneal epithelial cells.

### 3.3 Mdivi-1 preserves mitochondrial homeostasis and promotes corneal recovery following nitrogen mustard-induced injury in vivo

#### 3.3.1 Mdivi-1 promotes corneal repair and preserves tissue integrity following nitrogen mustard-induced injury

To determine whether the mitochondrial protective effects of Mdivi-1 observed in vitro translate into therapeutic benefit in vivo, we evaluated its efficacy in a previously established mouse model of acute mustard keratopathy [33]. Age-matched C57BL/6 mice (n=12 per group; 6 males and 6 females) were exposed to 40 mM nitrogen mustard on the ocular surface and treated with vehicle, 0.1% dexamethasone, or 50 μM Mdivi-1 beginning 2h after injury and continuing twice daily for 14 days. Dexamethasone served as positive control, representing the current anti-inflammatory approach for managing nitrogen mustard-induced ocular injury. In contrast, Mdivi-1 targets an upstream pathogenic mechanism by inhibiting DNM1L-mediated mitochondrial fission and preserving mitochondrial homeostasis.

Corneal epithelial integrity was monitored longitudinally by fluorescein staining over a 28-day period (Fig. 10A). Vehicle-treated eyes exhibited persistent corneal epithelial injury characterized by extensive fluorescein uptake and delayed recovery throughout the study period. In contrast, both dexamethasone and Mdivi-1 treated eyes showed marked reductions in fluorescein staining beginning at day 7, with progressive improvement through days 14, 21, and 28. Quantification of normalized corneal fluorescein staining scores demonstrated significant improvement in epithelial barrier function in both treatment groups compared with vehicle-treated controls (Fig. 10B). While vehicle-treated eyes achieved only 21% recovery by day 28, dexamethasone and Mdivi-1 treated eyes exhibited 86% and 92% improvement, respectively (Fig. 10C). Although the difference between dexamethasone and Mdivi-1 was not statistically significant, Mdivi-1 consistently demonstrated a trend toward greater efficacy throughout the recovery period.

**Figure 10.**
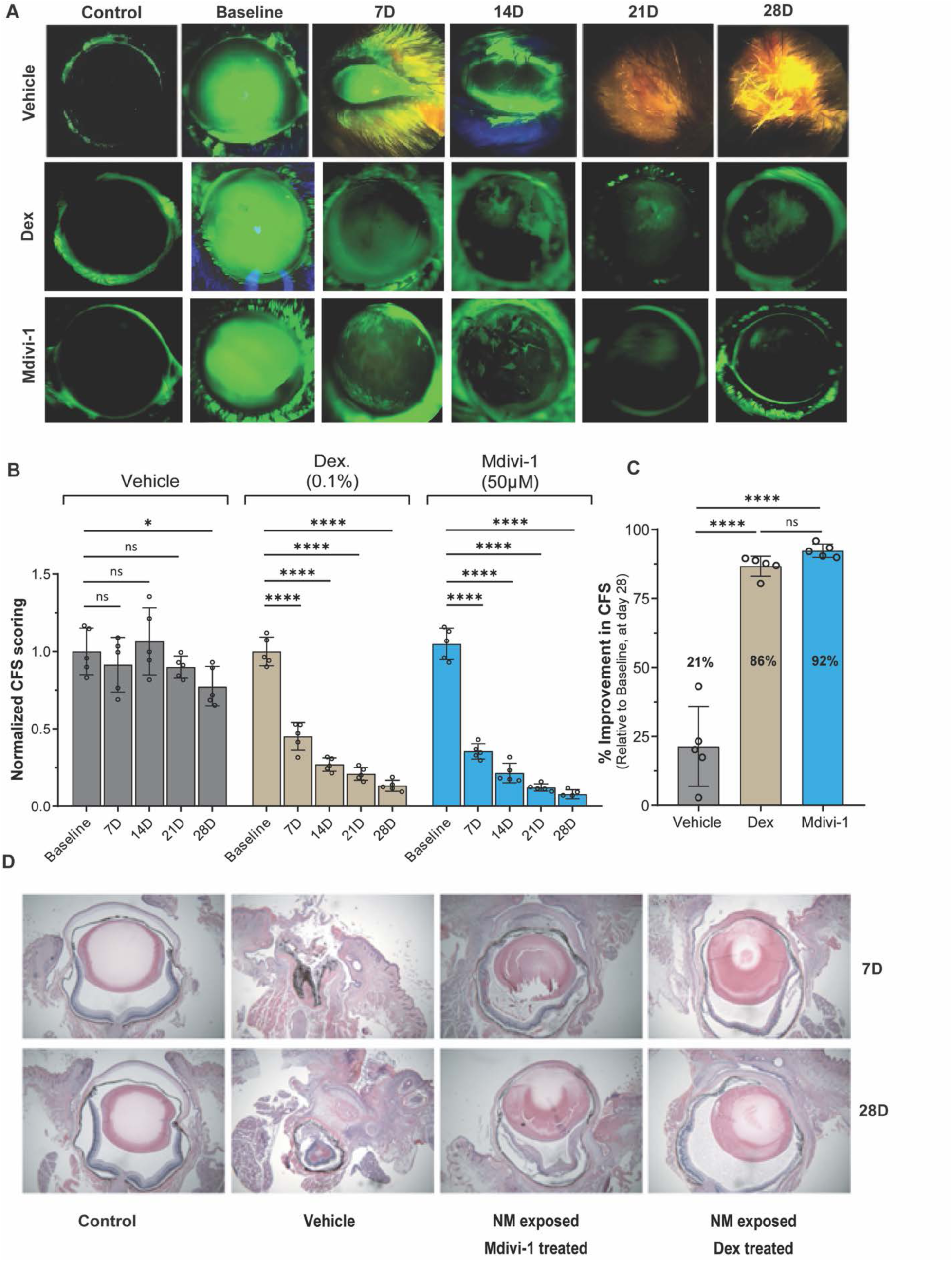
Mdivi-1 promotes corneal repair and preserves tissue integrity following nitrogen mustard-induced ocular injury in vivo. Age-matched C57BL/6 mice (n = 12 per group; 6 males and 6 females) were exposed to 40 mM nitrogen mustard and treated with vehicle, 0.1% dexamethasone, or 50 μM Mdivi-1 beginning 2 h after injury. (A) Representative fluorescein-stained corneas showing corneal epithelial damage and recovery at baseline and days 7, 14, 21, and 28 following nitrogen mustard exposure. Both dexamethasone and Mdivi-1 reduced fluorescein uptake compared with vehicle-treated eyes. (B) Quantification of normalized corneal fluorescein staining (CFS) scores over time. (C) Percent improvement in CFS at day 28 relative to baseline injury. Vehicle-treated eyes showed 21% recovery, whereas dexamethasone- and Mdivi-1-treated eyes exhibited 86% and 92% improvement, respectively. (D) Representative H&E-stained corneal sections collected at days 7 and 28 following nitrogen mustard exposure. Vehicle-treated corneas displayed epithelial erosion, stromal disorganization, and severe tissue damage, whereas Mdivi-1- and dexamethasone-treated corneas showed preservation of epithelial integrity and corneal architecture. Data are presented as mean ± SEM. *P < 0.05, ****P < 0.0001.

To determine whether improved corneal surface healing was accompanied by preservation of tissue architecture, histological analyses were performed at days 7 and 28 following nitrogen mustard exposure (Fig. 10D). Vehicle-treated corneas displayed severe tissue destruction, including epithelial erosion, stromal disorganization, and marked disruption of normal corneal architecture. In contrast, both dexamethasone and Mdivi-1-treated corneas exhibited substantial preservation of tissue structure, with improved epithelial continuity and maintenance of stromal organization. Notably, Mdivi-1-treated corneas demonstrated preservation of corneal architecture comparable to dexamethasone and showed a more intact epithelial surface relative to vehicle-treated controls.

These findings indicate that 50 µM Mdivi-1, delivered 2 hours after exposure to nitrogen mustard, promotes corneal epithelial repair and preserves ocular tissue integrity following nitrogen mustard-induced injury.

#### 3.3.2 Optimization of nitrogen mustard injury severity for therapeutic evaluation in mustard keratopathy

While topical application of nitrogen mustard reliably induced severe ocular surface injury, the extent of tissue damage often progressed rapidly and could obscure detection of therapeutic effects. Therefore, to establish a more controlled and reproducible model suitable for evaluating mitochondrial-targeted interventions, we adopted a localized nitrogen mustard exposure paradigm in which a 1.5-mm filter paper disc saturated with nitrogen mustard solution was applied to the central cornea for 2.5 min. This approach restricted injury primarily to the central cornea and enabled titration of injury severity while preserving surrounding tissue.

To identify an exposure level suitable for therapeutic studies, corneas were exposed to increasing concentrations of nitrogen mustard (500 μM, 5 mM, 20 mM, and 40 mM) and monitored longitudinally by fluorescein staining (Fig. S8A). Whereas 500 μM nitrogen mustard produced only mild injury and substantial spontaneous recovery, 40 mM nitrogen mustard caused severe ocular inflammation, extensive corneal damage, and progressive eye closure, limiting longitudinal assessment and therapeutic evaluation. Among the concentrations tested, 20 mM nitrogen mustard produced robust and reproducible corneal epithelial injury while maintaining sufficient ocular integrity for follow-up analyses. Based on these findings, 20 mM nitrogen mustard was selected as the optimal concentration for subsequent studies.

Importantly, the 20 mM exposure paradigm generated consistent corneal pathology while preserving adequate tissue architecture for comprehensive downstream analyses, including fluorescein staining, histological evaluation, apoptosis assessment, mitochondrial dynamics, mitochondrial integrity, and corneal nerve analysis.

Using this optimized injury paradigm, we evaluated the therapeutic efficacy of 50 μM Mdivi-1. Representative fluorescein images demonstrated progressive recovery of the corneal surface in Mdivi-1-treated eyes compared with vehicle-treated controls (Fig. S8B). Quantification of normalized corneal fluorescein staining scores revealed significantly improved epithelial healing in the Mdivi-1 group, with differences becoming apparent by day 14 and persisting through day 28 (Fig. S8C). By day 28, Mdivi-1-treated eyes exhibited approximately 75% improvement relative to baseline injury, compared with 32% improvement in vehicle-treated controls (Fig. S8D). These findings establish 20 mM nitrogen mustard as a reproducible and experimentally tractable model of mustard keratopathy and further support the therapeutic efficacy of Mdivi-1 in promoting corneal repair following vesicant-induced injury.

#### 3.3.3 Mdivi-1 preserves mitochondrial quality by reducing oxidative stress and promoting mitochondrial turnover

Our next set of studies employed the new method of nitrogen mustard delivery at 20 mM, applied to MitoTimer reporter mice. This mouse line harbors the Mitotimer reporter gene, which colocalizes to mitochondria, making it possible to directly visualize mitochondrial health following nitrogen mustard injury in vivo. Newly synthesized and functionally active mitochondria fluoresce green, whereas aged, oxidized, or dysfunctional mitochondria fluoresce red (Fig. 11A).

**Figure 11.**
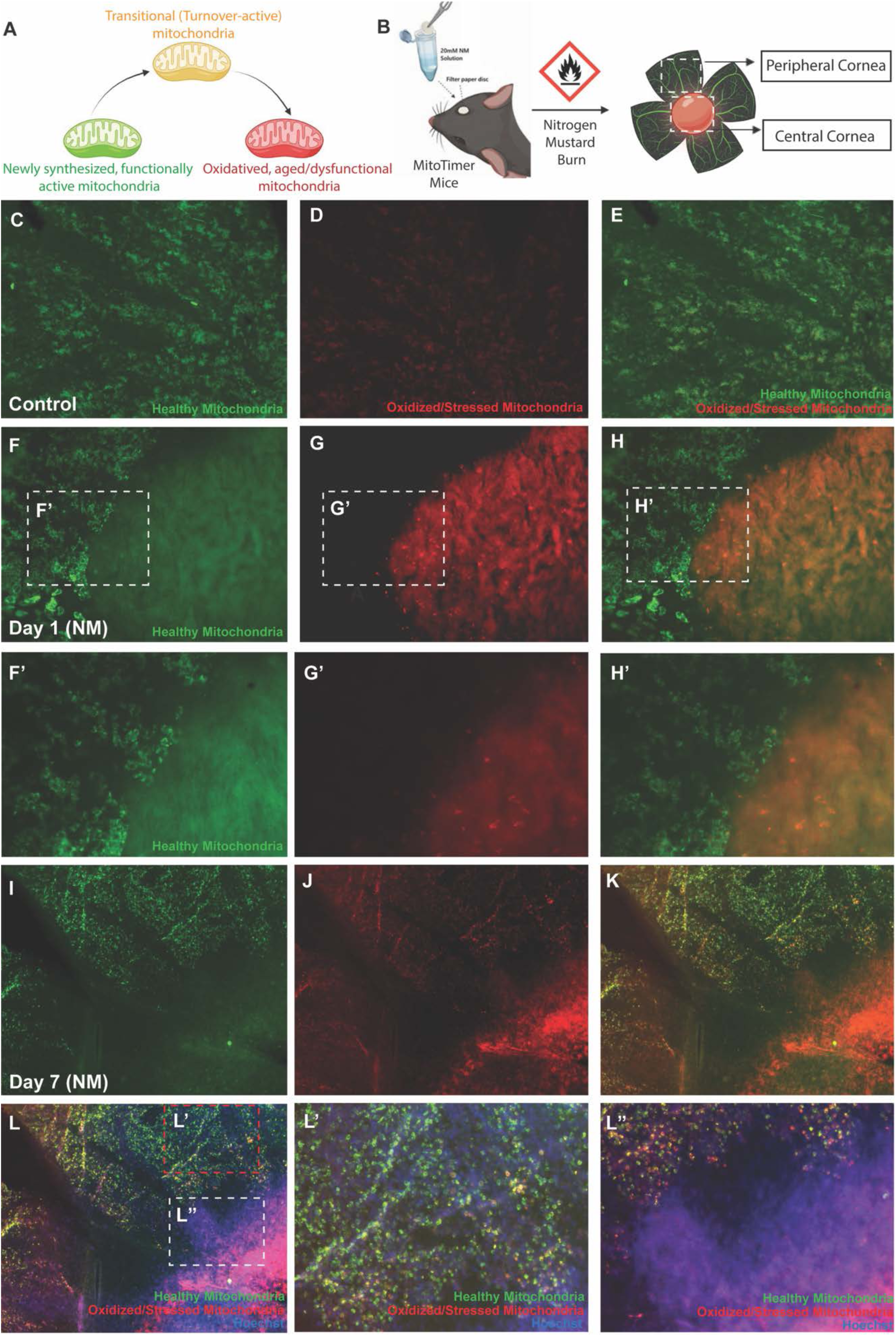
Nitrogen mustard exposure induces persistent mitochondrial oxidative stress and disruption of mitochondrial homeostasis in the cornea of MitoTimer mice. (A) Schematic of the MitoTimer reporter system. Newly synthesized and functionally active mitochondria fluoresce green, whereas aged, oxidized, or dysfunctional mitochondria fluoresce red. (B) Experimental design showing localized corneal exposure to 20 mM nitrogen mustard and assessment of mitochondrial status in central and peripheral corneal regions. (C–E) Representative corneal flat mounts from untreated control mice showing predominantly healthy mitochondrial populations with limited oxidized mitochondria. (F–H′) Representative corneal flat mounts one day after nitrogen mustard exposure demonstrating accumulation of oxidized/stressed mitochondria within the central cornea, the primary site of injury. Boxed regions are shown at higher magnification in F′–H′. (I–L″) Representative corneal flat mounts seven days after nitrogen mustard exposure showing persistent mitochondrial oxidative stress, expansion of oxidized mitochondrial populations, and disruption of mitochondrial homeostasis. Merged images include Hoechst nuclear staining (blue). Intense Hoechst staining within the central cornea is consistent with inflammatory cell infiltration at the injury site. Boxed regions are shown at higher magnification in L′ and L″.

We first verified MitoTimer reporter gene expression within the undamaged corneal epithelium. Corneal flat mounts were immunostained with cytokeratin 12 (K12), a corneal epithelial-specific marker. Both green and red MitoTimer fluorescence colocalized with K12-positive cells (Fig. S9), confirming that the MitoTimer signal originated predominantly from corneal epithelial cells. Importantly, flat-mount imaging enabled selective visualization of the corneal surface, facilitating assessment of epithelial mitochondrial dynamics while minimizing contributions from underlying stromal and other corneal cell populations.

Localized corneal injury was induced using the optimized 20 mM nitrogen mustard exposure model, and mitochondrial status was evaluated in corneal flat mounts at days 1 and 7 following injury (Fig. 11B). Under basal conditions, corneal epithelial cells exhibited predominantly green fluorescence with limited red signal, consistent with balanced mitochondrial turnover and maintenance of mitochondrial homeostasis (Fig. 11C–E). Notably, red fluorescence was modestly enriched within the central cornea relative to the periphery. This physiological gradient likely reflects normal corneal epithelial biology, as central epithelial cells are more differentiated, metabolically active, and farther removed from the limbal stem cell niche, resulting in reduced regenerative turnover and greater accumulation of older mitochondria. In contrast, the peripheral cornea, which is continuously replenished by limbal progenitor cells, exhibited predominantly green fluorescence indicative of a younger mitochondrial population.

As early as day 1 following nitrogen mustard exposure, a marked accumulation of red fluorescence was observed within the central cornea, corresponding to the primary site of injury, indicating rapid mitochondrial oxidative stress and damage (Fig. 11F–H′). By day 7, oxidized mitochondria remained elevated and extended throughout the injured region, demonstrating persistent disruption of mitochondrial homeostasis (Fig. 11I–L″). Notably, intense Hoechst staining was evident within the central cornea at day 7, consistent with substantial inflammatory cell infiltration at the site of injury. The spatial overlap between inflammatory cell accumulation and regions of elevated red fluorescence suggests a close association between persistent mitochondrial dysfunction and corneal inflammation following nitrogen mustard exposure. Together, these findings demonstrate that nitrogen mustard induces rapid and sustained mitochondrial oxidative damage that persists well beyond the initial injury.

We next examined whether Mdivi-1 preserves mitochondrial quality during the chronic phase of nitrogen mustard-induced corneal injury using corneal flat mounts collected 28 days after exposure (Fig. 12). In untreated control corneas, the mitochondrial population remained dominated by green fluorescence with minimal red signal, consistent with healthy mitochondrial turnover and homeostasis (Fig. 12A–C″). In contrast, nitrogen mustard-exposed vehicle-treated corneas exhibited persistent accumulation of red fluorescence throughout the injured cornea, particularly within the central region corresponding to the original site of injury (Fig. 12D–F″). Merged images demonstrated extensive yellow-orange fluorescence, indicating continued accumulation of oxidized and dysfunctional mitochondria despite four weeks of recovery. Notably, a distinct region within the central cornea exhibited near-complete loss of MitoTimer fluorescence (Fig. 12F, F′), suggesting severe cellular loss and tissue damage at the primary site of nitrogen mustard exposure.

**Figure 12.**
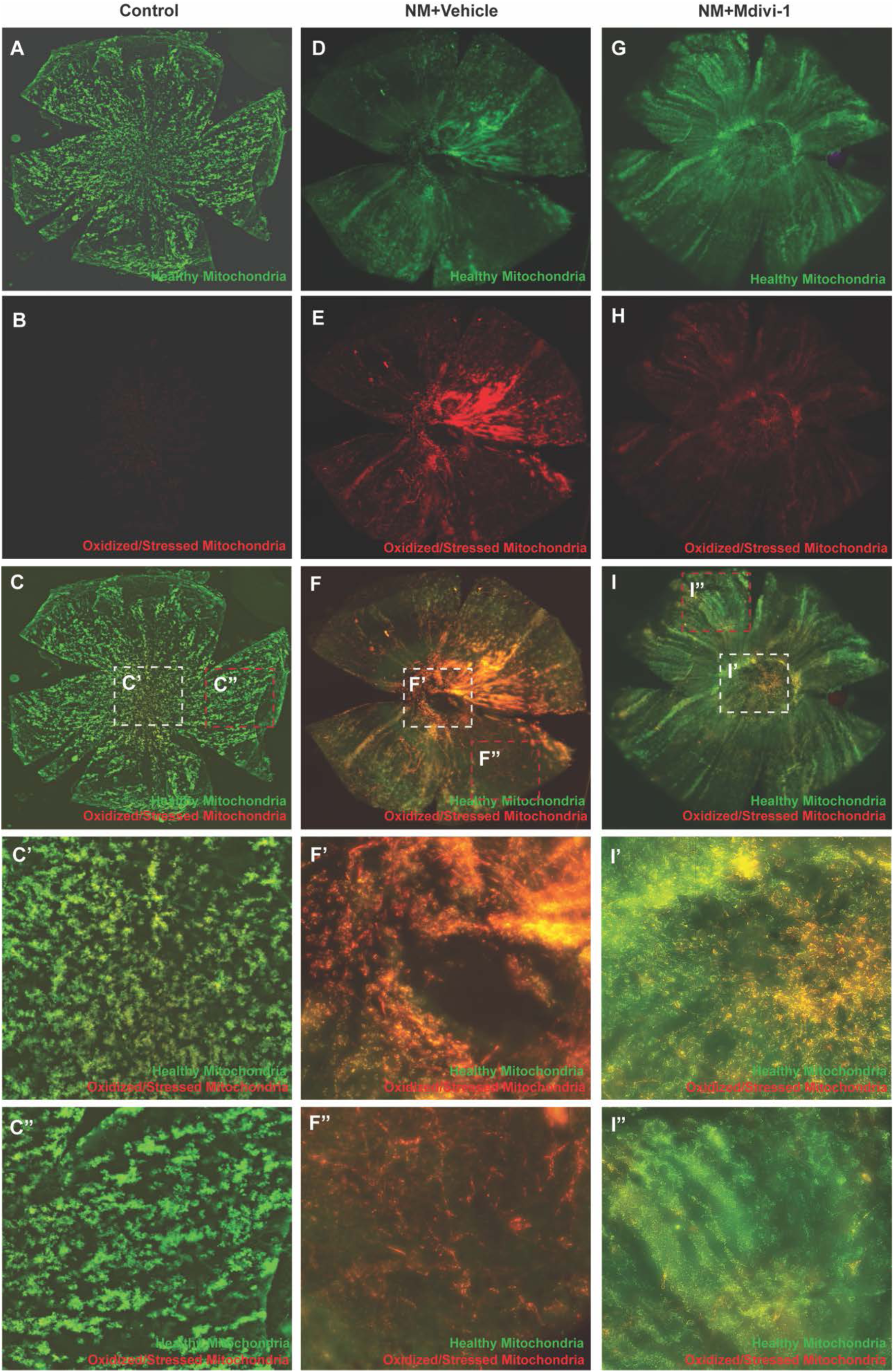
Mdivi-1 preserves mitochondrial quality and homeostasis following nitrogen mustard-induced corneal injury in MitoTimer mice. Corneal flat mounts collected 28 days after nitrogen mustard exposure showing healthy mitochondria (green), oxidized/stressed mitochondria (red), and merged images. (A–C) Untreated control corneas exhibiting predominantly healthy mitochondrial populations with minimal oxidized mitochondria. (D–F) NM-exposed vehicle-treated corneas displaying persistent accumulation of oxidized/stressed mitochondria, particularly within the central injury zone, resulting in extensive yellow-orange fluorescence in merged images. Notably, a focal region of near-complete loss of MitoTimer fluorescence is evident within the central cornea (F, F**′**), consistent with severe cellular loss and tissue damage at the primary site of nitrogen mustard exposure. (G–I) nitrogen mustard-exposed corneas treated with Mdivi-1 showing preservation of healthy mitochondrial populations and reduced accumulation of oxidized mitochondria. Higher-magnification images of the central cornea (**I′**) reveal predominantly green fluorescence with localized yellow fluorescence, indicative of active mitochondrial turnover and renewal. In the peripheral cornea (I″), green fluorescence is more prominent, consistent with maintenance of a healthy mitochondrial network and mitochondrial homeostasis. Boxed regions are shown at higher magnification in C′-C**″**, F′-F**″**, and I′-I**″**. Mdivi-1 treatment reduced mitochondrial oxidative stress, preserved mitochondrial organization, and restored a healthier mitochondrial profile throughout the injured cornea.

In contrast, Mdivi-1-treated corneas displayed substantially reduced red fluorescence and preservation of the healthy mitochondrial population (Fig. 12G–I″). An accumulation of mitochondria-rich cells was observed surrounding the injury site, suggesting active cellular repopulation and tissue repair within the damaged region. Higher-magnification images of the central cornea revealed predominantly green fluorescence together with localized yellow fluorescence (Fig. 12I′), indicative of healthy mitochondria undergoing active turnover and renewal. In the peripheral cornea, green fluorescence was even more pronounced (Fig. 12I″), consistent with maintenance of a healthy mitochondrial network and effective mitochondrial homeostasis. Compared with vehicle-treated corneas, Mdivi-1-treated corneas exhibited reduced mitochondrial oxidation, preservation of mitochondrial organization, and restoration of a more physiological mitochondrial profile across the ocular surface.

Collectively, these findings demonstrate that nitrogen mustard induces persistent mitochondrial dysfunction, inflammatory injury, and focal cellular loss that remain evident long after the initial exposure. Mdivi-1 preserves mitochondrial quality and homeostasis by limiting the accumulation of oxidized mitochondria, promoting active mitochondrial turnover, and supporting repopulation of the injured corneal epithelium during tissue repair.

## 4.0 Discussion

Mustard keratopathy remains a major unmet clinical challenge because currently available therapies primarily suppress inflammation [34, 35] without targeting the fundamental basis of tissue injury. Increasing evidence indicates that mitochondrial dysfunction is an early and sustained event in chemically induced ocular injury [36–39] contributing to oxidative stress, epithelial cell death, metabolic failure, and defective wound healing. Our study identifies excessive DNM1L-dependent mitochondrial fission as a central upstream event linking nitrogen mustard exposure to these pathological processes. Nitrogen mustard exposure caused rapid mitochondrial fragmentation was accompanied by membrane depolarization, mitochondrial reactive oxygen species accumulation, impaired antioxidant defenses, defective mitochondrial quality control and metabolic reprogramming. Pharmacological inhibition of mitochondrial fission with Mdivi-1 effectively interrupted this pathogenic cascade both in vitro and in vivo, establishing preservation of mitochondrial homeostasis as a promising therapeutic strategy for the treatment of mustard keratopathy.

Mitochondrial dynamics have emerged as critical regulators of cellular adaptation to stress [40–42], and excessive DNM1L activation is increasingly recognized as a common mechanism underlying neurodegenerative and ocular diseases [43–46]. Previous studies show that DNM1L-mediated mitochondrial fragmentation drives respiratory dysfunction, reactive oxygen species production, and cell death [47–50], whereas its inhibition preserves mitochondrial integrity, protects retinal pigment epithelial cells from oxidative damage, suppresses choroidal neovascularization, and improves tissue survival [51–54]. Consistent with these observations, nitrogen mustard rapidly increased DNM1L phosphorylation at Ser616 and disrupted the mitochondrial network in our study, while both Mdivi-1 and the selective DNM1L inhibitor Drp1i27 prevented mitochondrial fragmentation and preserved mitochondrial function. The ability of Drp1i27 to phenocopy Mdivi-1 is consistent with the idea that excessive DNM1L-dependent mitochondrial fission is a driver of nitrogen mustard-induced mitochondrial injury.

Oxidative stress is a hallmark of mustard-induced ocular injury [3, 7, 55–57], yet the cellular origin of persistent reactive oxygen species generation has remained incompletely defined. Previous studies have largely attributed vesicant toxicity to DNA alkylation, glutathione depletion, and inflammatory reactive oxygen species production [4, 56, 58–64]. Our findings expand this model by identifying mitochondrial fragmentation as a major upstream amplifier of oxidative injury. Importantly, Mdivi-1 exhibited minimal direct antioxidant activity in a cell-free assay but markedly reduced mitochondrial reactive oxygen species in intact cells while restoring endogenous antioxidant defenses. These findings suggest that the protective effects of Mdivi-1 arise primarily from preservation of mitochondrial function rather than direct radical scavenging, highlighting the therapeutic advantage of targeting the source of oxidative stress instead of its downstream consequences.

A particularly important finding of this study is the restoration of mitochondrial quality control following inhibition of mitochondrial fission. Mitochondrial quality control, through coordinated mitophagy and mitochondrial biogenesis, is essential for eliminating dysfunctional mitochondria and maintaining metabolic competence. Defective mitochondrial turnover has recently been implicated in chronic inflammation, impaired epithelial repair, and progressive tissue degeneration across multiple diseases [65–68]. Consistent with these reports, nitrogen mustard exposure impaired mitophagy, reduced mitochondrial DNA content, and promoted accumulation of oxidized mitochondria, whereas Mdivi-1 restored mitochondrial turnover, enhanced mitochondrial biogenesis, and significantly improved respiratory function. These observations are consistent with the idea that inhibition of mitochondrial fission not only prevents mitochondrial injury but actively promotes mitochondrial renewal, thereby re-establishing mitochondrial homeostasis. However, additional experiments will be needed to prove these processes. Future studies will include flux-based mitophagy experiments, and markers such as LC3-II, p62, PINK1, and Parkin, as well as assessment of mitochondrial biogenesis markers, including PGC-1α.

Restoration of mitochondrial quality translated directly into improved cellular bioenergetics. Nitrogen mustard profoundly suppressed oxidative phosphorylation and shifted ATP production toward glycolysis, reflecting severe mitochondrial dysfunction. Mdivi-1 significantly restored basal respiration, maximal respiration, spare respiratory capacity, and mitochondrial ATP production while normalizing intracellular pH, demonstrating recovery of mitochondrial metabolic competence. These findings reinforce the concept that mitochondrial dynamics regulate not only organelle morphology but also cellular energy metabolism and stress adaptation, consistent with recent advances in mitochondrial biology.

The mitochondrial protection observed in vitro translated into robust therapeutic efficacy in vivo. Topical Mdivi-1 accelerated corneal epithelial regeneration and maintained mitochondrial quality following nitrogen mustard exposure. Despite the severity of the injury model, Mdivi-1 demonstrated therapeutic efficacy comparable to, and in several outcome measures modestly greater than, dexamethasone. Unlike corticosteroids, which primarily suppress downstream inflammatory responses, Mdivi-1 targets an upstream pathogenic mechanism that preserves mitochondrial integrity, cellular metabolism, and tissue repair. These findings raise the possibility that mitochondrial-directed therapies could complement or reduce dependence on corticosteroids or other anti-inflammatory agents, while providing broader protection against vesicant-induced ocular injury.

Mdivi-1 has generated much interest as a potential therapeutic in numerous degenerative diseases that involve mitochondrial dysfunction and has been shown to be protective in several preclinical disease models including heart/brain ischemia-reperfusion injury [69, 70], traumatic brain injury [71], and Parkinson’s disease [72], corneal alkali burn [38], glaucoma [73, 74] and other optic neuropathies [75]. Although originally identified as a selective inhibitor of DNM1L-mediated mitochondrial fission, subsequent studies reported that Mdivi-1 exposure can also reversibly inhibit mitochondrial complex I and alter cellular bioenergetics [76, 77]. These biological effects appear to be highly context-dependent, varying with cell type, disease model, treatment duration, and dosing regimen. In the present study, corneal epithelial cells were exposed to Mdivi-1 for only 2 h in vitro, while mice received topical administration of 10 μL of 50 μM Mdivi-1 twice daily. Under these experimental conditions, we observed no evidence of mitochondrial toxicity or impaired cellular function. Instead, Mdivi-1 consistently preserved mitochondrial membrane potential, reduced mitochondrial ROS production, restored mitochondrial quality control and oxidative phosphorylation, and significantly improved corneal epithelial repair both in vitro and in vivo. Furthermore, the selective DNM1L inhibitor Drp1i27 reproduced the protective effects of Mdivi-1, providing independent pharmacological support consistent with the idea that inhibition of excessive DNM1L-dependent mitochondrial fission is one mechanism underlying the therapeutic benefit observed.

## 5.0 Conclusion

Collectively, our findings provide support for the hypothesis that excessive DNM1L-dependent mitochondrial fission is a central pathogenic mechanism driving nitrogen mustard-induced corneal injury. By preserving mitochondrial homeostasis, limiting oxidative stress, restoring mitochondrial quality control, and improving bioenergetic function, Mdivi-1 may promote corneal repair through a mechanism fundamentally distinct from conventional anti-inflammatory therapy. These findings suggest targeting mitochondrial dynamics may be an effective therapeutic strategy for mustard keratopathy. Mdivi-1 analogues are currently under development for human use, providing an opportunity for future therapeutic application.

## 6.0 Limitations of the study

Several limitations should be acknowledged. First, HCLE is an immortalized human limbal epithelial cell and cannot fully represent primary human corneal epithelium, limbal stem cells, or the corneal stroma. We plan to conduct studies with both primary human corneal epithelial cell lines and cultured corneal biopsy “buttons” in a future study.

Second, the 40 mM nitrogen mustard model produced an exceptionally severe ocular injury, characterized by extensive inflammation and, in some animals, profound structural damage, including loss of ocular globe integrity. Although the therapeutic activity of Mdivi-1 under these stringent conditions supports the robustness of its protective effects, the severity of the injury may have limited detection of its full therapeutic benefit and complicated assessment of corneal epithelial healing.

Third, the mechanistic conclusions were based primarily on pharmacological inhibition. However, the ability of the selective DNM1L inhibitor Drp1i27 to reproduce the major protective effects of Mdivi-1 substantially strengthens the evidence that excessive DNM1L-dependent mitochondrial fission contributes to nitrogen mustard-induced corneal injury. Future studies incorporating genetic modulation of DNM1L will provide additional confirmation and help define the specific cellular pathways involved.

Finally, longer-term studies will be important for determining whether inhibition of mitochondrial fission prevents recurrent epithelial breakdown, stromal fibrosis, neovascularization, chronic inflammation, corneal nerve degeneration, and visual dysfunction. Further investigation of the upstream signaling events responsible for nitrogen mustard -induced DNM1L activation will also clarify how vesicant exposure initiates mitochondrial fragmentation and may reveal additional therapeutic targets.

## CRediT authorship contribution statement

- Conceptualization: AGM, MEF
- Data curation: AGM, MEF
- Formal analysis: AGM, EM
- Funding acquisition: MEF
- Investigation: AGM, EM
- Methodology: AGM, HA, ZY, CDW, MEF
- Project administration: MEF
- Resources: MTC, ZY
- Supervision: MEF
- Visualization: AGM
- Writing – original draft: AGM, MEF
- Writing – review and editing: AGM, EM, HA, RR, MTC, CDW, MEF

## Declaration of generative AI use

The authors did not use any generative AI tools in preparing or revising the manuscript.

## Acknowledgments

The authors gratefully acknowledge Ilene K. Gipson, PhD, Schepens Eye Research Institute, Massachusetts Eye and Ear, Harvard Medical School for the immortalized human corneal limbal epithelial (HCLE) cell line and Tat Fong Ng, Department of Ophthalmology, Boston University School of Medicine for expert histological work.

The authors are grateful to Mei Zhan and Tiffany Draper of Virginia Tech for assistance in the transferring the MitoTimer expression construct and MitoTimer Reporter mice to Tufts University.

The authors further acknowledge Sharmila Masli, PhD, Boston University School of Medicine, Darlene Dartt, PhD, Schepens Eye Research Institute, Massachusetts Eye and Ear, Harvard Medical School and Jia Yin, MD, PhD, Schepens Eye Research Institute, Massachusetts Eye and Ear, Harvard Medical School for advice and support during the course of this study.

## Data and materials availability

All data supporting the findings of this study are available within the paper and its Supplementary Information. The MitoTimer reporter construct and MitoTimer transgenic reporter mice are available from Dr. Zhen Yan upon written request.

