## supplemental tables and figures for "Targeting Mitochondrial Dysfunction with Mdivi-1 Confers Therapeutic Protection in Cell Culture and Mouse Models of Mustard Keratopathy"

### Supplemental Material

**Figure S1. Dose-response optimization of DRP1i27 following nitrogen mustard exposure.** HCLE cells were exposed to 200 µM nitrogen mustard and treated with increasing concentrations of DRP1i27 (5-100 µM). Cell viability was assessed by WST-1 absorbance at 440 nm. Maximal protection was observed at 50 µM, which was selected for subsequent experiments. Data are presented as mean ± SEM.

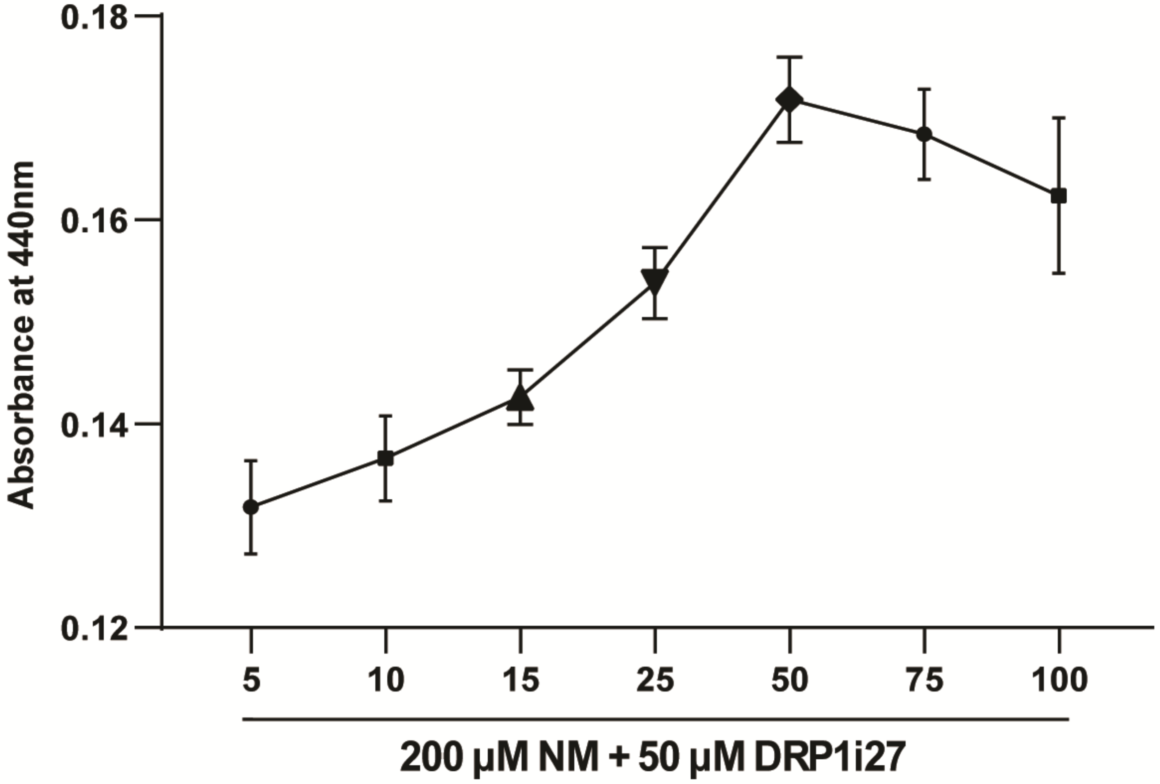

**Figure S2. DRP1i27 preserves mitochondrial morphology following nitrogen mustard exposure.** HCLE cells were stained with MitoTracker Green (green) and Hoechst (blue). Representative images show control cells (A-C), 200 μM nitrogen mustard -treated cells (D-F), and cells treated with 200 μM nitrogen mustard plus 50 μM DRP1i27 (G-I). DRP1i27 preserved mitochondrial morphology following nitrogen mustard exposure, consistent with the therapeutic effects observed with Mdivi-1. Boxed regions are enlarged in C′, F′, and I′.

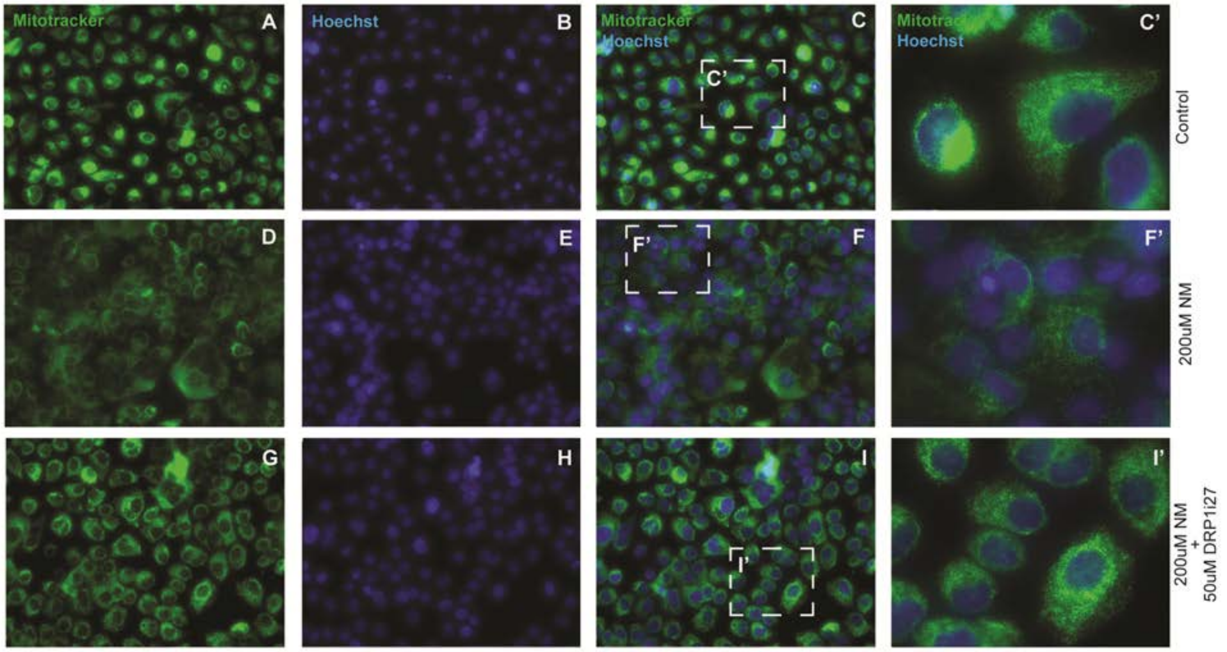

**Figure S3. DRP1i27 preserves mitochondrial membrane potential following nitrogen mustard exposure.** Mitochondrial membrane potential was assessed by JC-1 staining. Red JC-1 aggregates indicate polarized mitochondria, whereas green monomers indicate mitochondrial depolarization. Representative images show control cells (A-C), 200 μM nitrogen mustard -treated cells (D-F), and cells treated with 200 μM nitrogen mustard plus 50 μM DRP1i27 (G-I). DRP1i27 preserved mitochondrial membrane potential following nitrogen mustard exposure, consistent with the therapeutic effects obtained with Mdivi-1 and supporting inhibition of DRP1-dependent mitochondrial fission as the protective mechanism.

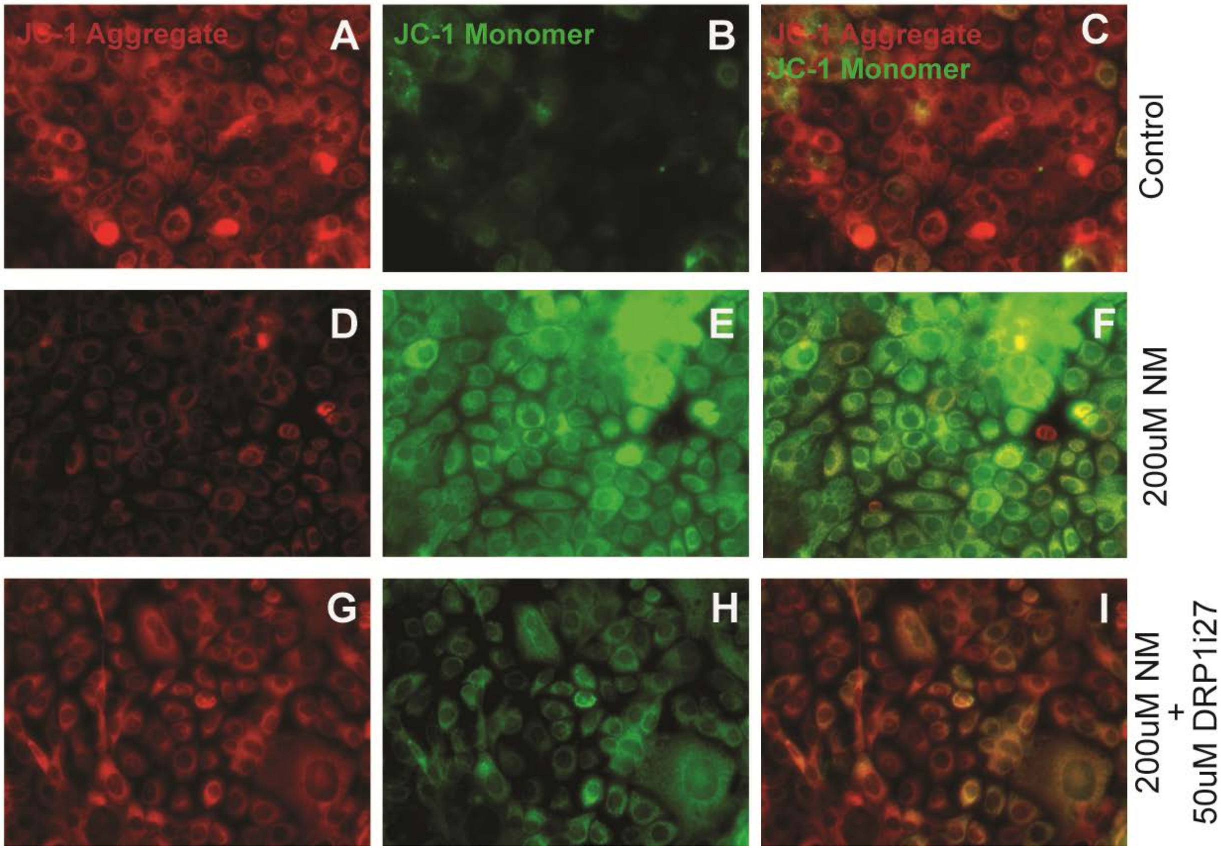

**Figure S4. DRP1i27 reduces nitrogen mustard-induced mitochondrial superoxide production.** Mitochondrial superoxide was assessed using MitoSOX Red, with Hoechst nuclear counterstaining. Representative images show control cells (A-C), 200 μM nitrogen mustard -treated cells (D-F), and cells treated with 200 μM nitrogen mustard plus 50 μM DRP1i27 (G–I). DRP1i27 reduced nitrogen mustard -induced mitochondrial superoxide, consistent with the antioxidant and mitochondrial protective effects observed with Mdivi-1.

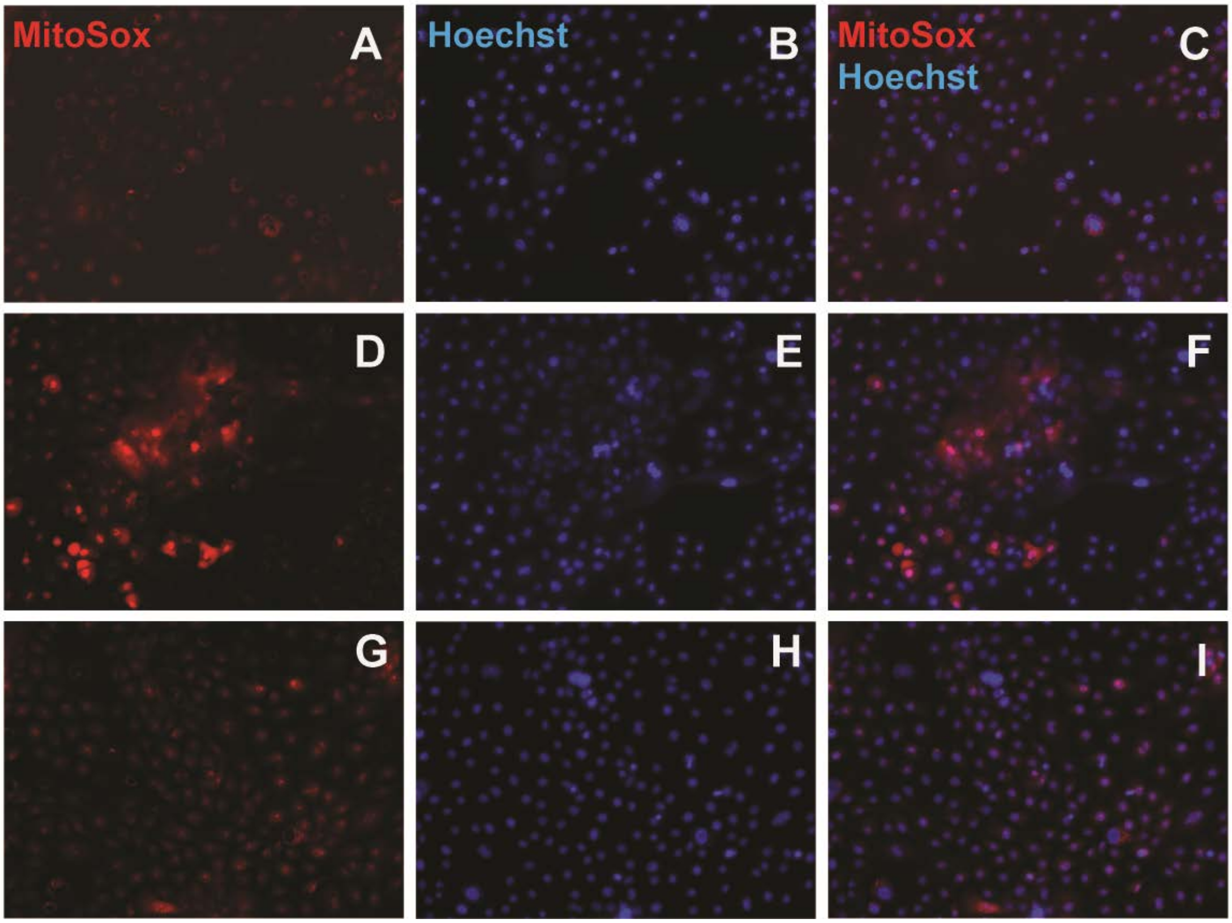

**Figure S5. Mdivi-1 restores catalase and eNOS expression following nitrogen mustard exposure.** HCLE cells were exposed to 200 μM nitrogen mustard, with or without 50 μM Mdivi-1. Catalase (A) and eNOS (B) mRNA expression was measured by qPCR, normalized to GAPDH, and expressed as $\log_2$ fold change relative to untreated controls. *P < 0.05, **P < 0.01, ***P < 0.001.

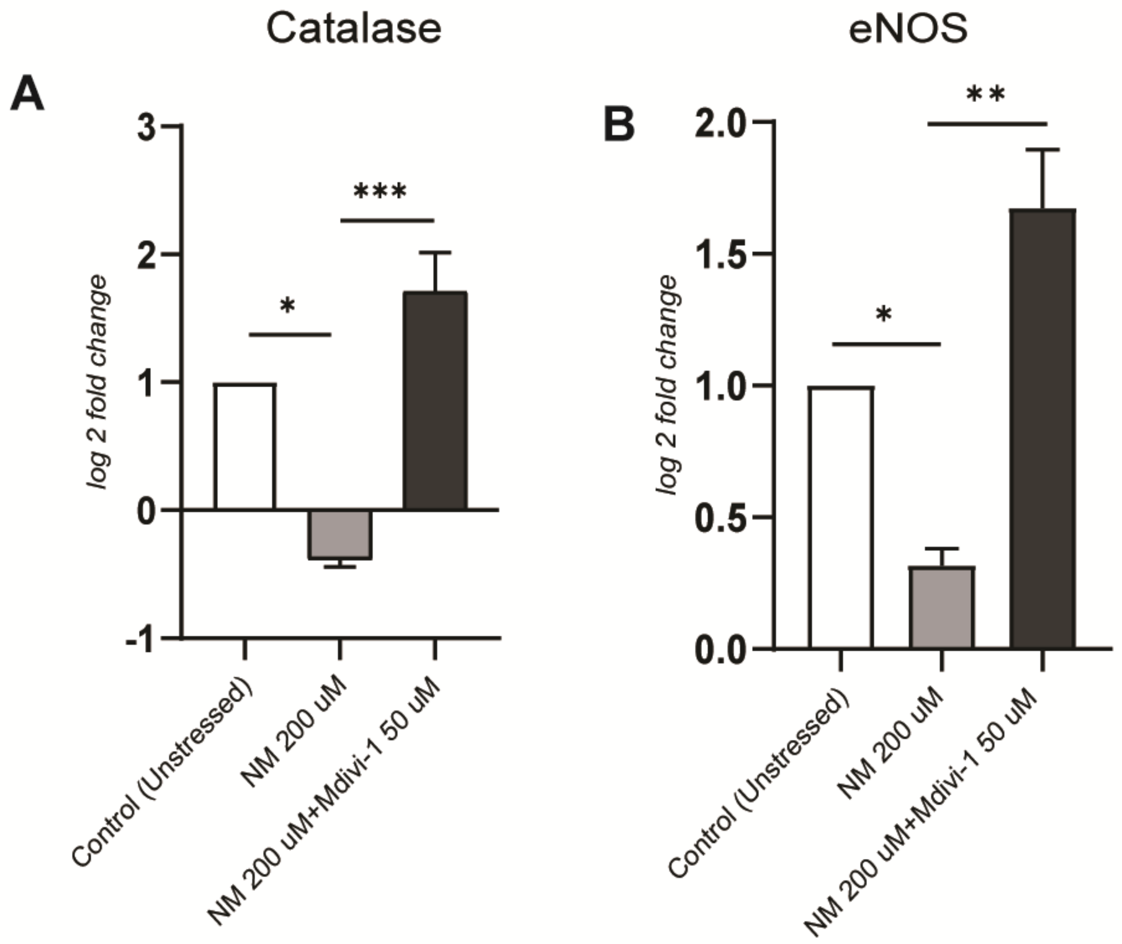

**Figure S6. Cell-free assessment of the antioxidant capacity of Mdivi-1.** Total antioxidant capacity was measured at 490 nm across increasing concentrations of Mdivi-1. Uric acid and glucose served as positive and negative controls, respectively. Mdivi-1 produced only a weak concentration-dependent signal compared with uric acid.

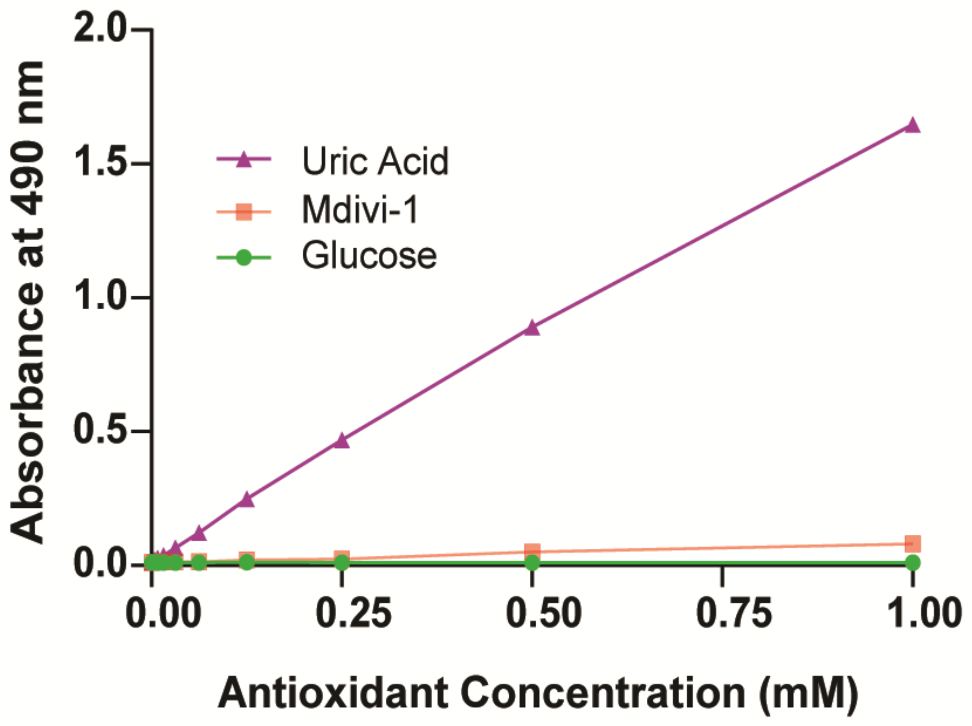

**Figure S7. Mdivi-1 prevents nitrogen mustard-induced loss of mitochondrial copy number in HCLE cells.** Mitochondrial biogenesis was assessed by quantifying mitochondrial DNA (mtDNA) copy number relative to nuclear DNA (nDNA). Mdivi-1 significantly preserved mitochondrial copy number at levels higher than those observed in nitrogen mustard-treated cells, consistent with maintenance and replenishment of the mitochondrial pool. Data are presented as mean ± SEM. *P < 0.05.

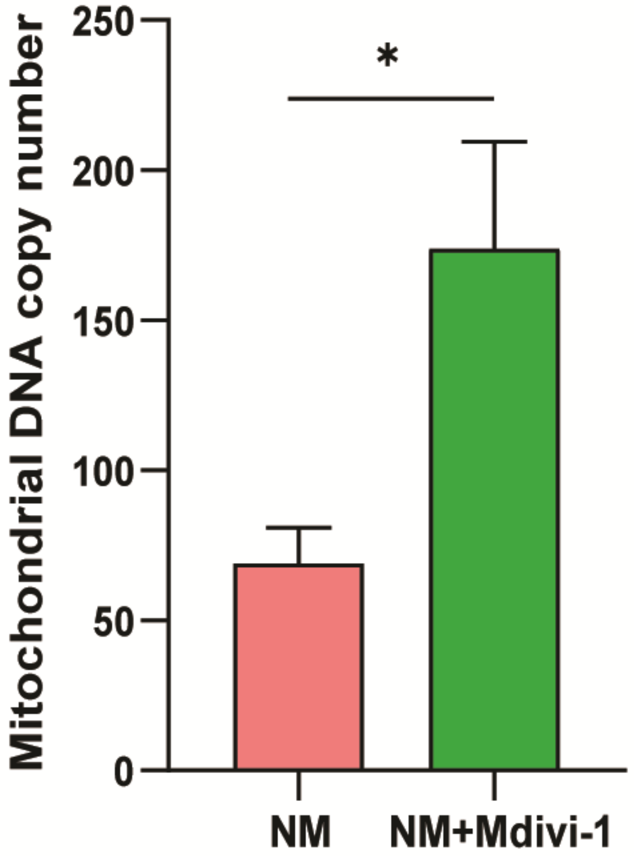

**Figure S8. Optimization of nitrogen mustard injury severity and validation of Mdivi-1 efficacy in a localized model of mustard keratopathy.** (A) Representative fluorescein-stained corneas following localized exposure to increasing concentrations of nitrogen mustard (500 μM, 5 mM, 20 mM, and 40 mM) using a central corneal application paradigm. Corneal injury was monitored at baseline and days 7, 14, 21, and 28. Injury severity increased in a concentration-dependent manner, with 40 mM nitrogen mustard producing severe inflammation and progressive eye closure, whereas 20 mM nitrogen mustard generated reproducible corneal injury while preserving ocular integrity for longitudinal assessment. (B) Representative fluorescein-stained corneas from vehicle- and Mdivi-1-treated mice following exposure to 20 mM nitrogen mustard. Mdivi-1 treatment promoted progressive restoration of the corneal epithelial surface over the 28-day observation period. (C) Quantification of normalized corneal fluorescein staining (CFS) scores following 20 mM nitrogen mustard exposure. Mdivi-1 significantly accelerated epithelial recovery compared with vehicle-treated controls. (D) Percent improvement in CFS at day 28 relative to baseline injury. Mdivi-1-treated eyes exhibited 75% improvement compared with 32% improvement in vehicle-treated controls. Data are presented as mean ± SEM. *P < 0.05, **P < 0.01, ***P < 0.001, ****P < 0.0001.

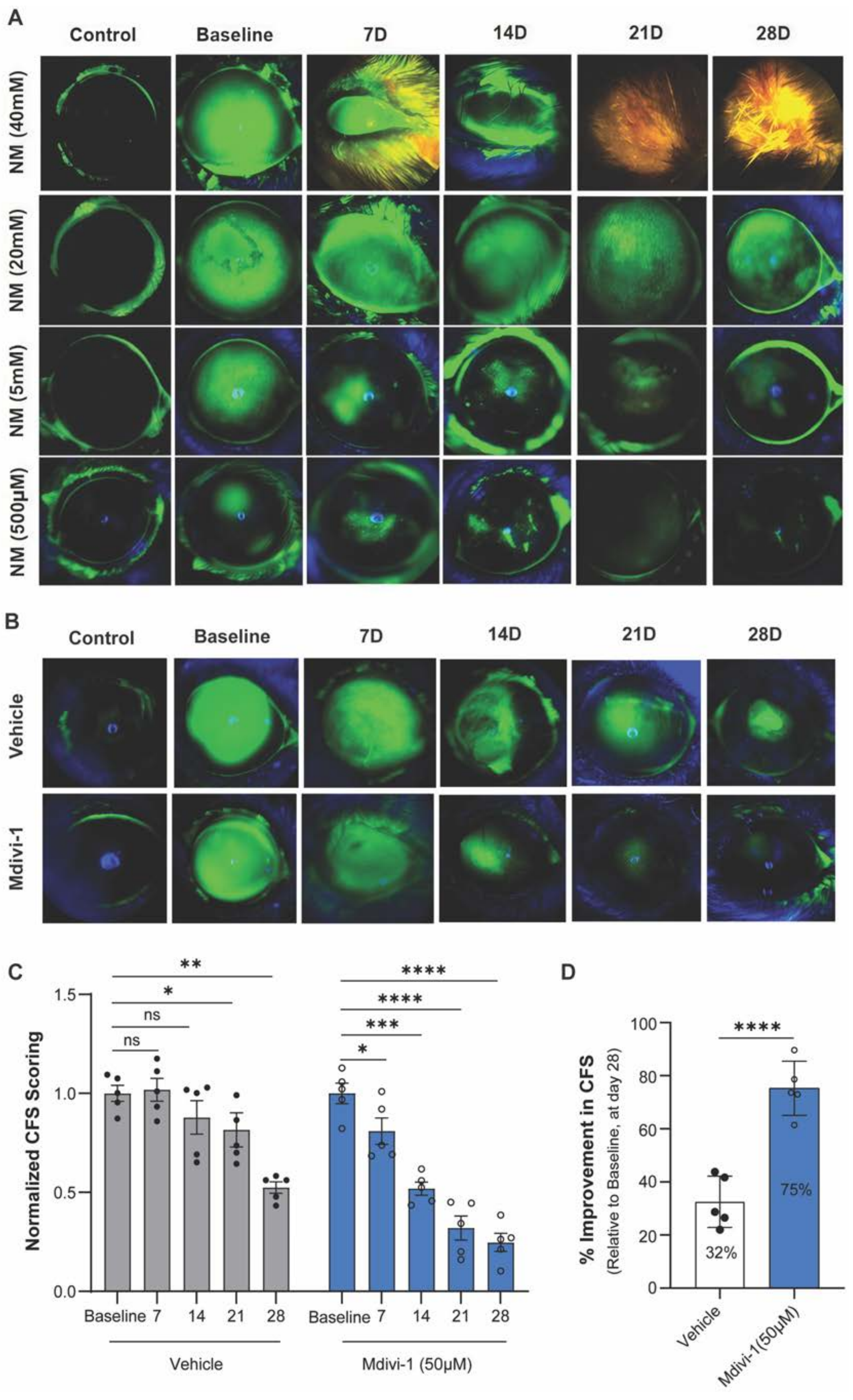
A
Control
Baseline
7D
14D
21D
28D
NM (40mM)
NM (20mM)
NM (5mM)
NM (500µM)
B
Control
Baseline
7D
14D
21D
28D
Vehicle
Mdivi-1
C
Normalized CFS Scoring
1.5
1.0
0.5
0.0
**
*
ns
ns
****
****
***
*
Baseline 7 14 21 28
Baseline 7 14 21 28
Vehicle
Mdivi-1 (50µM)
D
% Improvement in CFS
(Relative to Baseline, at day 28)
100
80
60
40
20
0
****
32%
75%
Vehicle
Mdivi-1(50µM)

**Figure S9. Validation of corneal epithelial-specific MitoTimer expression in MitoTimer mice.** Representative corneal cross-sections from healthy control MitoTimer reporter mice immunostained for cytokeratin 12 (CK12), a marker of differentiated corneal epithelial cells. (A) Newly synthesized/healthy mitochondria (green). (B) Oxidized/stressed mitochondria (red). (C) CK12 immunostaining (cyan). (D) Overlay of oxidized/stressed mitochondria with K12. (E) Overlay of healthy mitochondria with K12. (F) Merged image showing colocalization of both mitochondrial populations with CK12-positive corneal epithelial cells. These images confirm that MitoTimer fluorescence is predominantly localized to the corneal epithelium, supporting its use for assessing epithelial mitochondrial dynamics and quality control in vivo.

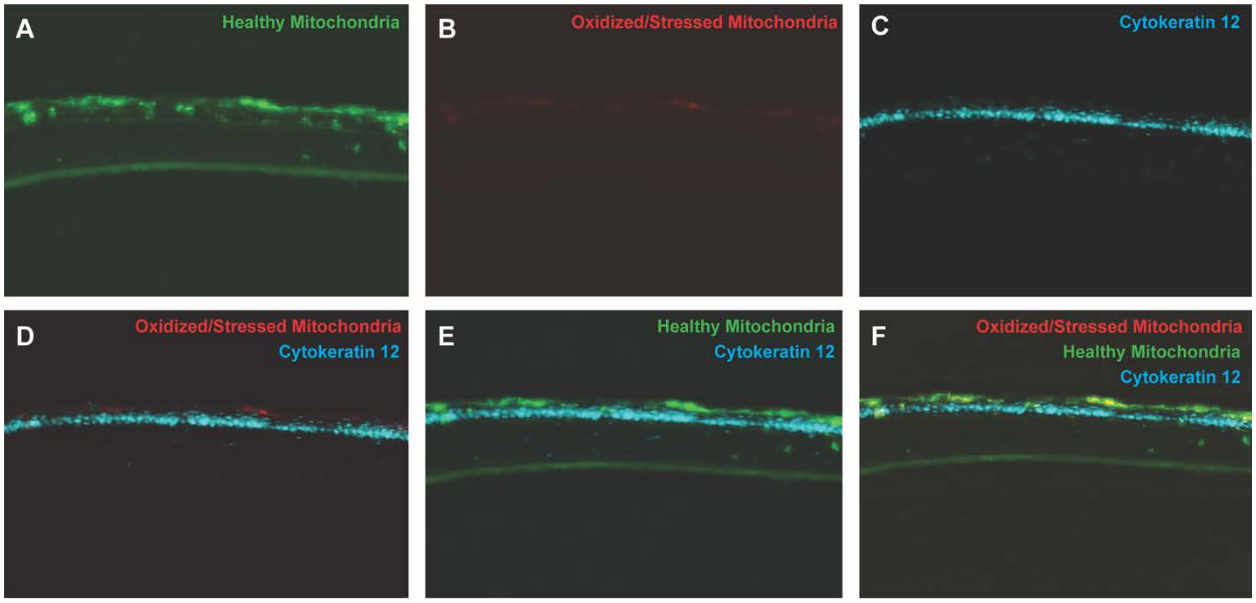
